# Multi-layered regulatory networks driving human dopaminergic neuron differentiation

**DOI:** 10.64898/2026.06.30.735499

**Authors:** Dicle Malaymar Pinar, Yaxin Jing, Haonan Li, Xiaonan Liu, Andrea Coschiera, Juha Kere, Masahito Yoshihara, Peter Swoboda, Pelin Sahlén, Markku Varjosalo

**Affiliations:** Institute of Biotechnology, HiLIFE, University of Helsinki, Helsinki, Finland; Department of Medicine Huddinge (MedH), Karolinska Institutet, Stockholm, Sweden; Institute for Advanced Academic Research, Chiba University, Chiba, Japan; Department of Artificial Intelligence Medicine, Chiba University Graduate School of Medicine, Chiba, Japan; Department of Advanced Artificial Intelligence Medicine, The University of Osaka Graduate School of Medicine, Suita, Osaka, Japan; Science for Life Laboratory, School of Engineering Sciences in Chemistry, Biotechnology and Health, KTH Royal Institute of Technology, Solna, Sweden; Folkhälsan Research Centre, Helsinki, Finland; Stem Cells and Metabolism Research Program, University of Helsinki, Helsinki, Finland; iCAN Digital Precision Cancer Medicine Flagship, University of Helsinki, Helsinki, Finland

## Abstract

Neuronal differentiation requires coordinated regulation across chromatin organization, gene expression, protein abundance, and post-translational modifications. Using the LUHMES human dopaminergic neuronal differentiation model, we integrated proteome and phosphoproteome profiling with previously generated enhancer–promoter interaction maps from NET-CAGE and HiCap and transcriptomic analysis across three consecutive differentiation stages. Differentiation was accompanied by increased abundance and phosphorylation of proteins involved in axon guidance, cytoskeletal organization, and synaptic signaling, alongside repression of the cell cycle, DNA replication, and chromatin-associated programs. Phosphoproteome analysis further revealed extensive remodeling of signaling networks associated with neuronal maturation. Enhancer–promoter interaction analysis revealed substantially greater rewiring at enhancers than promoters and identified master and relay transcription factors regulated across multiple molecular layers. siRNA-mediated knockdown showed that transcription factors MYT1, ISL2, and NHLH2 are crucial for proper neuronal maturation, whereas LCOR acts as a negative regulator of differentiation. Integrative network reconstruction further nominated MEOX2 as a candidate enhancer-associated regulator of late dopaminergic maturation. Together, these findings provide a multi-layered view of regulatory networks governing human neuronal differentiation.

## INTRODUCTION

Neuronal differentiation is a multistep process in which progenitor cells gradually acquire the molecular and functional characteristics of mature neurons(Ming & Song, 2011). This process begins with the commitment of multipotent stem or progenitor cells to a neural lineage, followed by a series of tightly regulated molecular and cellular changes. These changes include exit from the cell cycle, morphological transformations such as neurite outgrowth, and the establishment of functional properties like synapse formation and neurotransmitter expression(Götz & Huttner, 2005; Wamsley & Fishell, 2017). Chromatin modifications, transcription factors, and signaling pathways coordinate these events to ensure proper neuronal identity and function(Yao & Jin, 2014; Nord & West, 2020). During differentiation, dynamic changes occur at multiple regulatory levels, including enhancer and promoter activation, transcriptional shifts, and alterations in protein expression(Feng *et al*, 2007; Ziller *et al*, 2015). These layers of regulation work together to precisely coordinate neuronal maturation and specialization, ultimately leading to the formation of functional neural circuits. Recent advances in multi-omics technologies have enabled integrated characterization of transcriptional, proteomic, and phosphoproteomic regulation during neuronal development and neurodegenerative diseases(Seddighi *et al*, 2024; Hao *et al*, 2026). Single-cell and spatial multi-omics studies have further demonstrated coordinated changes in chromatin accessibility, gene expression, and protein abundance during neural development(Guo *et al*, 2025; Uzquiano *et al*, 2022). Furthermore, three-dimensional chromatin architecture profiling using technologies such as Hi-C-based approaches has shown that enhancer–promoter interactions undergo extensive remodeling during neuronal differentiation, directly influencing transcriptional programs(Yoshihara *et al*, 2025; Heffel *et al*, 2024).

Understanding the molecular mechanisms underlying human neuronal development provides important insights into nervous system biology and neurodegenerative disease. Due to the limited accessibility of developing human neurons, in vitro models are widely used to investigate neuronal lineage commitment and maturation. LUHMES (Lund Human Mesencephalic) cells are a human neuronal precursor cell line that undergoes rapid and synchronized differentiation into post-mitotic dopaminergic-like neurons(Scholz *et al*, 2011). Upon tetracycline-induced suppression of the v-myc transgene, LUHMES cells exit the cell cycle, acquire neuronal morphology, and express neuronal markers including βIII-tubulin, MAP2, and tyrosine hydroxylase(Scholz *et al*, 2011; Lauter *et al*, 2020; Tüshaus *et al*, 2021). Importantly, compared with other neuronal cell systems, LUHMES cells retain a cytologically normal human karyotype (46 chromosomes without additional chromosomal abnormalities) and exhibit a highly synchronized and complete differentiation process, with no proliferating precursor cells remaining after the initiation of differentiation(Lauter *et al*, 2020). Proteomic and transcriptomic studies have established LUHMES cells as a robust human dopaminergic model system, showing strong induction of neuronal markers such as L1CAM, α-synuclein, MAPT, and SYN116, while recapitulating key aspects of human fetal midbrain development(Lauter *et al*, 2020; Tüshaus *et al*, 2021; Beliakov *et al*, 2023; Keighron *et al*, 2024). Compared with alternative neuronal models such as SH-SY5Y cell line, LUHMES cells also show high reproducibility and responsiveness to neurotoxic compounds(Lotharius *et al*, 2005; Beliakov *et al*, 2023; Keighron *et al*, 2024), making them well suited for systems-level studies of neuronal differentiation.

To investigate regulatory changes during neuronal differentiation, we conducted an integrative, multi-omics analysis of LUHMES cells collected at Day 1, Day 3, and Day 6 of differentiation. We combined enhancer-promoter interaction mapping using NET-CAGE(Hirabayashi *et al*, 2019) and HiCap(Su *et al*, 2021) (promoter-cHi-C) with transcriptomics, total proteomics, and phosphoproteomics to characterize coordinated changes in chromatin interactions, gene expression, protein abundance, and phosphorylation dynamics. NET-CAGE enables sensitive detection of nascent RNAs, including enhancer-derived transcripts, whereas HiCap identifies promoter–cis-regulatory element interactions at high resolution(Hirabayashi *et al*, 2019; Su *et al*, 2021). This integrated approach allowed us to connect chromatin-associated regulatory changes with downstream transcriptional and proteomic remodeling. As post-translational modifications are central regulators of neuronal signaling and development(Ochoa *et al*, 2020; Dumrongprechachan *et al*, 2021; Gay *et al*), phosphoproteome profiling further enabled characterization of signaling network remodeling during neuronal maturation.

## RESULTS

### Proteome profiling across LUHMES differentiation stages reveals dynamic and stage-specific protein expression

To characterize global proteome and phosphoproteome remodeling during LUHMES neuronal differentiation, we performed DIA-based mass spectrometry analysis on cells collected at three key stages: Day 1 (initiation), Day 3 (early differentiation), and Day 6 (maturation) (Fig. 1A). Across biological triplicates, approximately 7,600 proteins were identified at each stage (Fig. 1B). A total of 7,237 proteins were detected across all three time points, indicating broad preservation of the core proteome throughout differentiation (Fig. 1B). In contrast, subsets of proteins showed stage-specific expression, including 46 proteins uniquely detected at Day 1 and 112 uniquely detected at Day 6. Pairwise overlap analysis further identified 196 proteins shared exclusively between Day 1 and Day 3 and 178 shared between Day 3 and Day 6, consistent with progressive proteomic remodeling during neuronal maturation (Fig. 1B).

**Figure 1.**
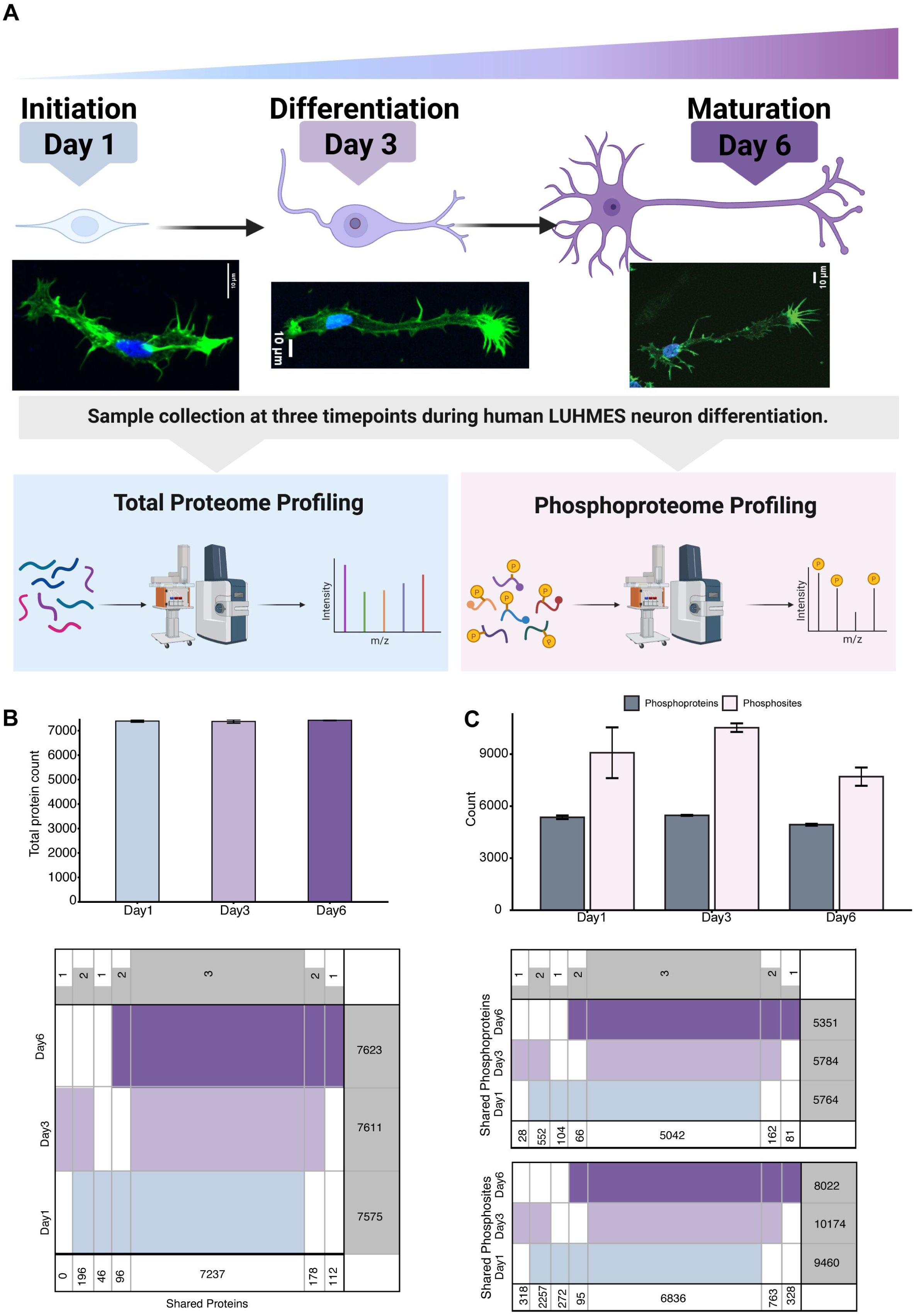
Overview of LUHMES cell differentiation and proteomic profiling strategy. **(A)** Schematic representation of the LUHMES neuronal differentiation timeline. Cells were collected at three stages: initiation (Day 1), differentiation (Day 3), and maturation (Day 6). Samples from each time point were processed for total proteome and phosphoproteome analysis by mass spectrometry. Representative fluorescence images are shown for each stage, with Hoechst-stained nuclei and phalloidin-stained F-actin indicating overall cell anatomy. Scale bars, 10 μm. **(B)** Bar plots showing the number of proteins identified in each biological replicate (n = 3) at Day 1, Day 3, and Day 6. A SuperVenn diagram summarizes the overlap of identified proteins across the three time points. **(C)** Bar plots showing the number of phosphoproteins and phosphosites identified in each biological replicate (n = 3) at each time point. SuperVenn diagrams summarize the overlap of phosphoproteins and phosphosites across Day 1, Day 3, and Day 6.

Phosphoproteome profiling identified approximately 5,000–5,800 phosphoproteins and 8,000–10,000 phosphosites per stage, with more than 6,800 phosphosites shared across all time points (Fig. 1C). Principal component analysis of both proteome and phosphoproteome datasets showed clear separation of Day 1, Day 3, and Day 6 samples along PC1, explaining 44.3% and 45.2% of total variance, respectively (Supplementary Fig. 1A,B). Hierarchical clustering similarly grouped samples according to differentiation stage, supporting the reproducibility and stage-specific organization of the datasets (Supplementary Fig. 1C,D).

### Differential protein expression dynamics during LUHMES neuronal differentiation

To assess stage-specific changes in protein abundance during LUHMES differentiation, we performed pairwise differential expression analyses for the Day 3 vs Day 1, Day 6 vs Day 3, and Day 6 vs Day 1 comparisons (Fig. 2A,C,E; Supplementary Table 1). Proteins with an adjusted p-value < 0.05 and |log₂(fold change)| > 1 were considered significantly regulated.

**Figure 2.**
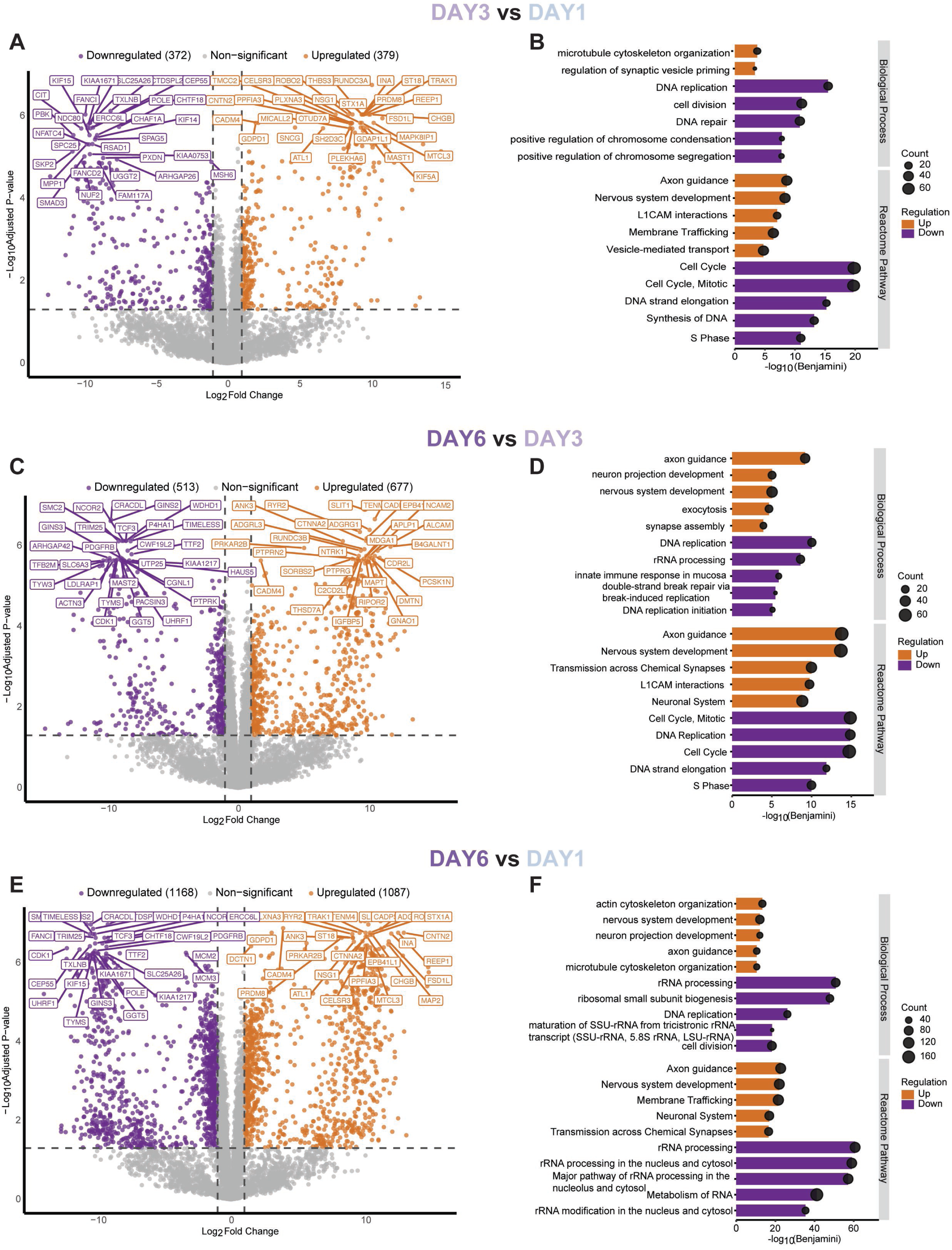
Differential protein abundance and functional enrichment analysis during LUHMES differentiation. Volcano plots show pairwise comparisons of protein abundance between Day 3 vs Day 1 **(A)**, Day 6 vs Day 3 **(C)**, and Day 6 vs Day 1 **(E)**. Proteins are plotted by log_2_ (fold change) (x-axis) and –log_10_ (adjusted p-value) (y-axis). Proteins with an adjusted p-value < 0.05 and |log₂(fold change)| > 1 were considered significantly differentially expressed. Significantly upregulated proteins are shown in orange, downregulated proteins in purple, and non-significant proteins in grey. The top 30 upregulated and downregulated proteins are labeled. Bar plots show Gene Ontology Biological Process and Reactome pathway enrichment analysis of differentially expressed proteins for Day 3 vs Day 1 **(B)**, Day 6 vs Day 3 **(D)**, and Day 6 vs Day 1 **(F)**. Only the top five enriched terms per regulation direction with a Benjamini-adjusted p-value < 0.001 are shown. Bar length represents statistical significance (–log₁₀ adjusted p-value), circle size indicates the number of proteins associated with each term, and upregulated and downregulated protein sets were analyzed separately. Complete enrichment results are provided in Supplementary Table 2.

The transition from Day 1 to Day 3 was characterized by balanced upregulation (379 proteins) and downregulation (372 proteins), reflecting early differentiation-associated remodeling (Fig. 2A). Downregulated proteins were predominantly associated with DNA replication and cell cycle progression including MCM complex components (MCM2, MCM4–MCM7) and DNA polymerase subunits (POLA1/2, POLD1, POLE, POLE2), reflecting suppression of proliferative programs during cell cycle exit(Aria & Yeeles, 2019; Nick McElhinny *et al*, 2008). In contrast, upregulated proteins were enriched for cytoskeletal organization and early neuronal features including β-tubulin isoforms (TUBB2A, TUBB3, TUBB4A) and microtubule-associated proteins such as MAP1B, MAP2, and MAP6(Conde & Cáceres, 2009; Tantry & Santhakumar, 2023).

Between Day 3 and Day 6, the number of differentially regulated proteins increased (677 upregulated vs 513 downregulated), indicating progressive acquisition of neuronal features (Fig. 2C). Continued repression of MCM complex members and cell cycle regulators, including CDK1 and CDK2, further supported sustained cell cycle exit. Concurrently, proteins involved in synaptic vesicle dynamics (e.g., SYN1, SYT1, CADPS), neuronal adhesion (e.g., NRXN1, NRXN2, NCAM1, NCAM2), axonal and dendritic structural maturation (e.g., NEFM, MAP2, MAP1B), and voltage-gated calcium signaling (e.g., CACNA1B) were prominently upregulated together with established neuronal differentiation markers including ELAVL4, DCLK1, MAPT(Lauter *et al*, 2020).

The comparison between Day 6 and Day 1 revealed the most extensive proteome remodeling, with 1,087 upregulated and 1,168 downregulated proteins (Fig. 2E). Downregulated proteins prominently included core DNA replication machinery (MCM2–7, GINS1–4, CDC45, POLA1–POLE2, PRIM1/2), cell cycle regulators (CDK1/2/4, CHEK1, MKI67, PCNA)(Martínez-Alonso & Malumbres, 2020; Juríková *et al*, 2016), chromatin assembly and remodeling factors (HELLS, CHAF1A/B, DNMT1)(Hoffmann & Schulz, 2021), and condensin complex subunits (SMC2, SMC4, NCAPD2, NCAPG, NCAPH)(Jackson & Bartek, 2009). In parallel, proliferative signaling molecules (NOTCH1/2, STAT1/3/5B, YAP1)(Lui *et al*, 2011) and metabolic enzymes supporting nucleotide and amino acid biosynthesis (DHFR, IMPDH1/2, SHMT1, GFPT1/2)(Saha *et al*, 2023) were also reduced.

Conversely, proteins associated with axon guidance and neuronal adhesion (DCC, ROBO2/3, SLIT1, L1CAM, CNTN1/2, NCAM1/2)(Kolodkin & Tessier-Lavigne, 2011), synaptic vesicle trafficking components (STX1A/B, STXBP1/5, RIMS1/3/4, UNC13A/B, BSN, SYN1) and voltage-gated ion channels (CACNA1B/H, KCNQ2, SCN3A/B, HCN4) were markedly increased(Südhof, 2013; Alexander *et al*, 2015). Cytoskeletal proteins (TUBB2A/B, TUBB3, TUBB4A, MAP1A/B, MAP2, NEFL, NEFM) and axonal transport kinesins (KIF5A/C, KIF21A/B) also accumulated. Increased abundance of neuronal identity regulators, including MYT1, ISL2, ELAVL3/4, NOVA1/2, and MECP2, together with neuronal signaling kinases such as CDK5, CAMK4, FYN, and BRSK1/2, further supported progressive neuronal maturation(Hirokawa *et al*, 2010; Puri *et al*, 2023; Chen *et al*, 2020; Cheung & Ip, 2012).

Several established neuronal proteins were among the most strongly upregulated proteins during differentiation, including DCX, MAP2, L1CAM, DCC, CNTN2, NEFM, and NRXN1 (Supplementary Fig. 2A). In addition, multiple highly induced proteins without prior neuronal annotation were identified, including TMSB10, RGMB, REEP1, TMEFF1, and IGFBPL1 (Supplementary Fig. 2B).

Together, these data demonstrate a coordinated proteomic reorganization from proliferative and biosynthetic programs toward structural, synaptic, and signaling modules characteristic of post-mitotic neurons during LUHMES differentiation. Consistent with these molecular changes, immunofluorescence imaging showed progressive morphological remodeling, with compact Day 1 cells extending elongated neurites at Day 3 and forming complex interconnected neuronal networks by Day 6 (Fig. 1A).

### Functional enrichment of differentially abundant proteins reveals stage-specific biological processes

To interpret the functional relevance of differentially abundant proteins, Gene Ontology (GO) biological process and Reactome pathway enrichment analyses were performed separately for upregulated and downregulated proteins across each comparison (Fig. 2B,D,F; Supplementary Table 2). Enrichment was based on proteins with an adjusted p-value < 0.001 (Benjamini–Hochberg correction).

In the Day 3 vs Day 1 comparison, upregulated proteins were enriched for cytoskeletal organization, axon guidance, and early neuronal development, whereas downregulated proteins were associated with DNA replication, cell cycle progression, and chromosome organization (Fig. 2B). Reactome pathway analysis similarly revealed early activation of neuronal system and membrane trafficking pathways together with repression of mitotic and DNA replication programs.

Between Day 6 and Day 3, enrichment patterns shifted further toward neuronal maturation, with upregulated proteins associated with synapse assembly, vesicle-mediated transport, and nervous system development, while downregulated proteins remained strongly linked to DNA replication, cell cycle progression, and RNA-associated processes (Fig. 2D).

The Day 6 vs Day 1 comparison showed the most pronounced functional divergence. Upregulated proteins were predominantly enriched for neuronal differentiation, axon guidance, cytoskeletal remodeling, and synaptic signaling pathways, whereas downregulated proteins were associated with proliferation, RNA metabolism, ribosome biogenesis, and chromatin-related processes (Fig. 2F).

Together, these enrichment analyses demonstrate a progressive functional transition during LUHMES differentiation characterized by activation of neuronal structural and synaptic programs alongside coordinated repression of proliferative and biosynthetic pathways.

### Protein expression trajectories across LUHMES neuronal differentiation

To investigate temporal protein abundance dynamics during LUHMES differentiation, proteins were clustered according to their expression profiles across Day 1, Day 3, and Day 6. This analysis identified nine distinct trajectory patterns, including continuous increase, continuous decrease, early increase, early decrease, late increase, late decrease, peak at Day 3, valley at Day 3, and a large stable cluster with minimal variation across time points (Supplementary Fig. 3; Supplementary Table 3). Notably, the stable cluster comprised most quantified proteins (5,460 proteins, including 233 transcription factors), whereas dynamic clusters represented smaller but functionally specialized subsets. Subsequent analyses focused on the six major monotonic trajectories that showed progressive increases or decreases during differentiation (Fig. 3A–F).

**Figure 3.**
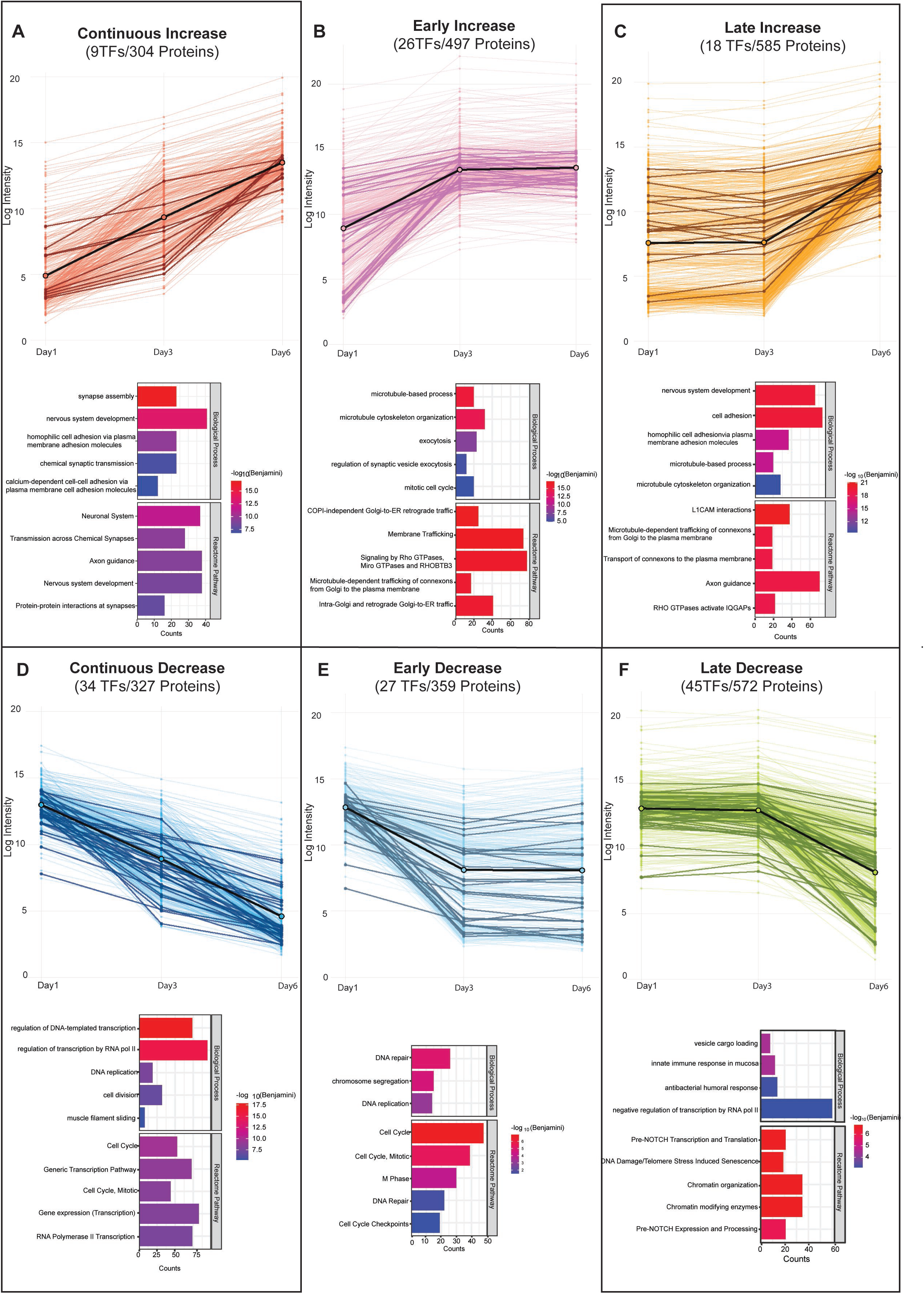
Clustering of protein expression trajectories during LUHMES differentiation. Line plots show protein abundance profiles across Day 1, Day 3, and Day 6, grouped into six clusters based on their temporal expression patterns: continuous increase **(A)**, early increase **(B)**, late increase **(C)**, continuous decrease **(D)**, early decrease **(E)**, and late decrease **(F)**. Each thin line represents an individual protein. Bold colored lines indicate transcription factors (TFs), and the bold black line represents the average expression profile of the cluster. The number of TFs and total proteins assigned to each cluster is indicated in each panel. Bar plots below each cluster show the top five enriched Gene Ontology Biological Process and Reactome pathway terms for proteins in the corresponding cluster. Bar length represents the number of proteins associated with each term, and color indicates statistical significance (–log₁₀ Benjamini-adjusted p-value).

Proteins in the continuous increase cluster (9 TFs / 304 proteins) showed gradual accumulation from Day 1 to Day 6 and were enriched for nervous system development, axon guidance, neuronal system pathways, and synapse assembly, consistent with progressive establishment of neuronal connectivity (Fig. 3A). Proteins in the early increase cluster (26 TFs / 497 proteins) increased primarily between Day 1 and Day 3 and were enriched for microtubule organization, exocytosis, membrane trafficking, and RHO GTPase signaling pathways, indicating early activation of cytoskeletal remodeling and vesicle transport programs (Fig. 3B). In contrast, proteins in the late increase cluster (18 TFs / 585 proteins) remained relatively stable during early differentiation but increased strongly at Day 6. These proteins were enriched for nervous system development, cell adhesion, axon guidance, and L1CAM interaction pathways, suggesting progressive activation of maturation-associated neuronal adhesion and connectivity programs (Fig. 3C).

Decreasing trajectory clusters were predominantly associated with proliferative and chromatin-related functions. Proteins in the continuous decrease cluster (34 TFs / 327 proteins) progressively declined throughout differentiation and were enriched for transcriptional regulation, DNA replication, and mitotic cell cycle pathways, consistent with sustained repression of proliferative programs (Fig. 3D). The early decrease cluster (27 TFs / 359 proteins) showed rapid reduction by Day 3 and was enriched for DNA replication, chromosome segregation, DNA repair, and cell cycle checkpoint pathways, reflecting early shutdown of mitotic machinery (Fig. 3E). Proteins in the late decrease cluster (45 TFs / 572 proteins) declined predominantly between Day 3 and Day 6 and were enriched for chromatin organization, chromatin-modifying enzymes, and NOTCH-related signaling pathways, indicating progressive attenuation of chromatin-associated regulatory programs during neuronal maturation (Fig. 3F).

### Phosphoproteome remodeling during LUHMES neuronal differentiation

To complement total proteome profiling, we quantified site-specific phosphorylation changes across LUHMES differentiation stages (Fig. 4A–F; Supplementary Table 4). Phosphosite intensities were analyzed without normalization to total protein abundance to preserve phosphorylation dynamics associated with large differentiation-dependent protein abundance changes. Given the large dynamic range of protein abundance changes across differentiation, normalization to total protein levels could reduce the apparent magnitude of phosphorylation changes on strongly upregulated neuronal proteins. Therefore, these data were interpreted as reflecting differentiation-associated phosphoproteome remodeling rather than phosphorylation occupancy. This interpretation is consistent with recent temporal proteomic and phosphoproteomic profiling of iPSC-derived neuronal differentiation, in which phosphoproteome dynamics were analyzed as a complementary regulatory layer to total proteome changes(Hao *et al*, 2026).

**Figure 4.**
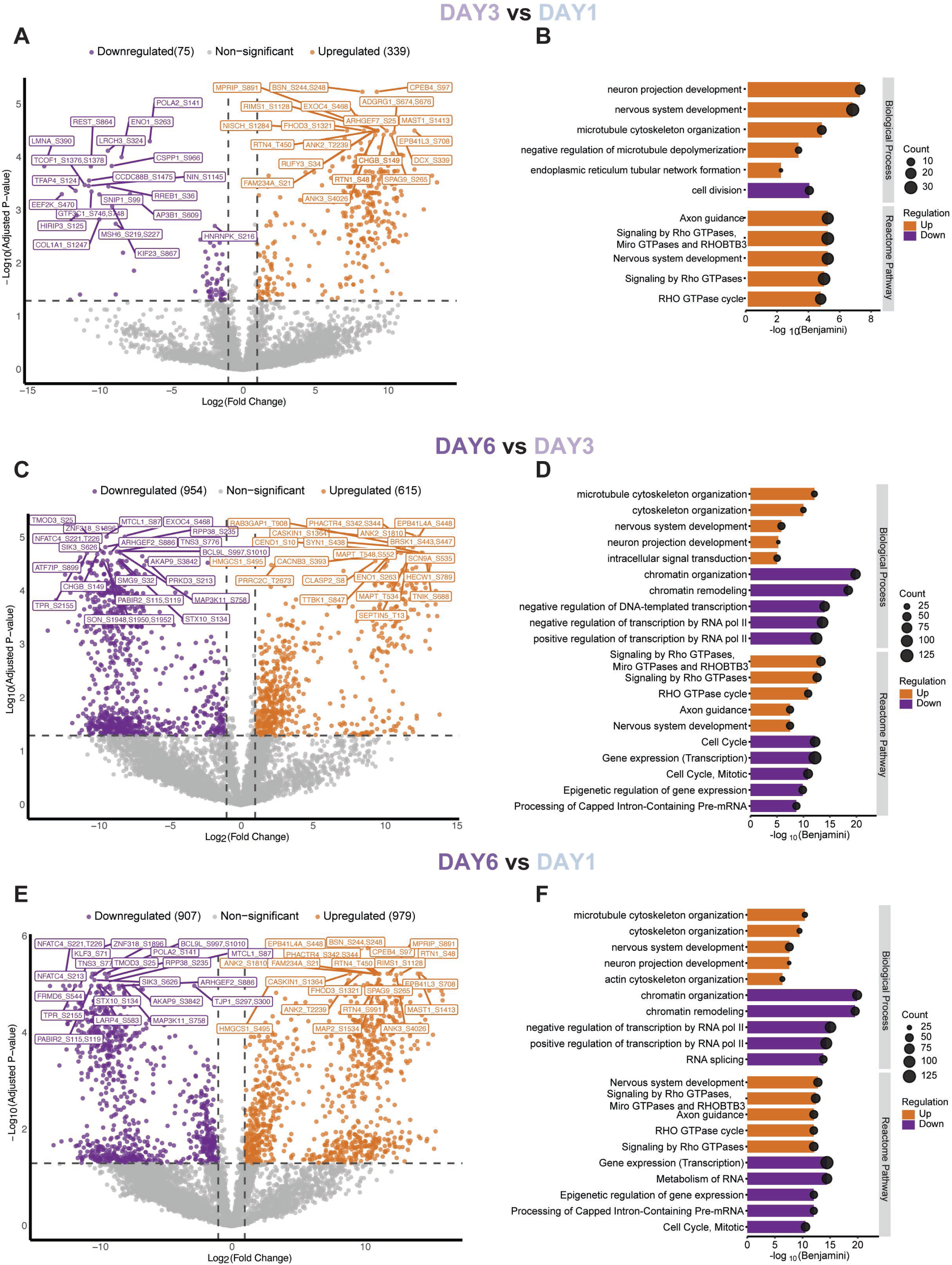
Differential phosphoproteome dynamics during LUHMES cell differentiation. Volcano plots show pairwise comparisons of phosphosite abundance between Day 3 vs Day 1 **(A)**, Day 6 vs Day 3 **(C)**, and Day 6 vs Day 1 **(E)**. Phosphosites are plotted by log_2_ (fold change) (x-axis) and –log_10_ (adjusted p-value) (y-axis). Phosphosites with an adjusted p-value < 0.05 and |log_2_(fold change)| > 1 were considered significantly regulated. Significantly increased phosphosites are shown in orange, decreased phosphosites in purple, and non-significant sites in grey. The top 20 increased and decreased phosphosites are labeled. Bar plots show Gene Ontology Biological Process and Reactome pathway enrichment analysis of proteins corresponding to significantly regulated phosphosites for Day 3 vs Day 1 **(B)**, Day 6 vs Day 3 **(D)**, and Day 6 vs Day 1 **(F)**. Bar length represents –log_10_(Benjamini-adjusted p-value), circle size indicates the number of proteins associated with each term and increased and decreased sets were analyzed separately.

The Day 3 vs Day 1 comparison identified 339 upregulated and 75 downregulated phosphosites, indicating early phosphoproteome remodeling during neuronal commitment (Fig. 4A). Downregulated phosphosites predominantly mapped to proteins associated with chromatin organization, DNA replication, and cell cycle regulation, including LMNA, EZH2, POLA2, LIG1, and MSH6. In contrast, increased phosphorylation predominantly affected proteins involved in cytoskeletal remodeling and neuronal morphogenesis, including ANK3, MAP2, MAP1B, DCX, BSN, RIMS1, and EXOC4. Increased phosphorylation was also observed on regulators of RHO-dependent cytoskeletal signaling and on the neuronal transcription factor MYT1, indicating extensive post-translational remodeling during early differentiation. Functional enrichment analysis identified cytoskeletal organization, neuron projection development, and RHO GTPase signaling pathways among upregulated phosphoproteins, whereas downregulated phosphoproteins were associated with proliferation-related processes (Fig. 4B; Supplementary Table 5).

Between Day 6 and Day 3, phosphoproteome remodeling became substantially more extensive, with 615 upregulated and 954 downregulated phosphosites (Fig. 4C). Reduced phosphorylation was observed on multiple transcriptional and chromatin-associated regulators, including ZBTB46, ZBTB18, NCOR2, SIN3A, CHD7, and SON, together with proteins involved in nuclear organization and intracellular trafficking. Conversely, increased phosphorylation was enriched among proteins associated with nervous system development, axon guidance, cytoskeletal remodeling, and RHO GTPase signaling pathways, whereas decreased phosphorylation remained linked to chromatin organization and transcriptional regulation pathways (Fig. 4D; Supplementary Table 5).

Direct comparison of Day 6 and Day 1 revealed extensive bidirectional phosphoproteome remodeling, with 979 upregulated and 907 downregulated phosphosites (Fig. 4E). Downregulated phosphosites prominently mapped to proteins associated with chromatin regulation, RNA processing, DNA replication, and genome maintenance, including NFATC4, SRRM2, NCOR2, LIG1, MDC1, and CHAF1A. In contrast, strongly increased phosphorylation was detected on neuronal cytoskeletal proteins and synaptic regulators, including NEFM, DCX, MAP2, MAPT, ANK2/3, L1CAM, RIMS1, and BSN, together with actin and RHO-associated cytoskeletal regulators such as MPRIP, FHOD3, and SPAG9. Functional enrichment analysis demonstrated strong enrichment of neuronal differentiation, cytoskeletal organization, axon guidance, and synaptic signaling pathways among upregulated phosphoproteins, whereas downregulated phosphoproteins were associated with chromatin remodeling, RNA metabolism, and cell cycle regulation (Fig. 4F; Supplementary Table 5).

To further investigate the functional consequences of phosphoproteome remodeling, we performed kinase–substrate enrichment analysis (KSEA) using the Day 6 vs Day 1 comparison (Supplementary Fig. 4; Supplementary Table 6). KSEA revealed a pronounced shift in kinase activity during differentiation, with activation of neuronal signaling kinases including DYRK2 and MAPK7 and suppression of multiple proliferation-associated kinases, including CDK1, CDK2, PLK1, and MAPKAPK2. Analysis of transcription factor substrates targeted by significantly regulated kinases identified coordinated phosphorylation changes across 67 TF substrates from 11 kinases, with substrates of suppressed CDKs and PLK1 showing widespread decreases in phosphorylation (Supplementary Fig. 4A,B; Supplementary Table 6).

Together, these findings demonstrate progressive phosphoproteome remodeling during LUHMES differentiation, characterized by extensive rewiring of kinase-driven signaling networks, coordinated suppression of proliferation- and chromatin-associated signaling, and activation of neuronal cytoskeletal, synaptic, and kinase signaling pathways.

### Remodeling of regulatory protein classes during LUHMES neuronal differentiation

We next examined changes in major regulatory protein classes during LUHMES differentiation, including kinases, phosphatases, transcription factors (TFs), small GTPases, and GPCRs (Supplementary Fig. 5A). In the Day 3 vs Day 1 comparison, transcription factors represented a substantial fraction of significantly regulated proteins, with 21 TFs upregulated and 23 downregulated. Kinases and phosphatases also showed pronounced changes at this stage. In the Day 6 vs Day 3 comparison, 17 TFs were upregulated and 48 were downregulated. The largest shift was observed in the Day 6 vs Day 1 comparison, where 33 TFs were upregulated and 85 were downregulated. Similar patterns were observed for kinases across all comparisons.

We next assessed the distribution of TF families among significantly regulated transcription factors (Supplementary Fig. 5B; Supplementary Table 1). Across all comparisons, C2H2 zinc finger proteins constituted the largest TF family, with particularly strong representation among downregulated TFs in the Day 6 vs Day 1 comparison. Additional TF families detected in both upregulated and downregulated groups included bHLH/bZIP, Homeodomain, HMG/Sox, EBF1/Ets, and SMAD/STAT factors.

We then integrated protein abundance and phosphorylation data for transcription factors significantly upregulated in the Day 6 vs Day 1 comparison (Supplementary Fig. 5C; Supplementary Table 1,4). Several TFs showed strong protein-level upregulation, including MYT1, ISL2, LHX2, DPF3, MEOX2, POU2F2, JUN, and NHLH2. Among these, BACH2,

MYT1, ST18, SCRT1, LCOR, ZNF316, and MECP2 also displayed significantly increased phosphorylation. Other upregulated TFs showed detectable but non-significant phosphorylation changes or lacked phosphoproteomic measurements.

To further characterize changes across regulatory enzyme classes, we analyzed differentially regulated enzymes and modifiers in the Day 6 vs Day 1 comparison (Supplementary Fig. 6A,B; Supplementary Table 7). Kinases and phosphatases represented the most abundant upregulated enzyme classes, whereas methyltransferases showed the strongest downregulation. Differential regulation was also observed across ubiquitin ligases, deubiquitinases, acetyltransferases, RNA-editing enzymes, and proteases. Volcano plot analysis revealed extensive remodeling of signaling-associated kinases together with chromatin- and RNA-associated regulatory enzymes (Supplementary Fig. 6B). Together, this class-level analysis provides a maturation-associated regulatory framework for identifying processes that become prominent as LUHMES cells transition from proliferative precursors to post-mitotic neurons. The coordinated increase in kinases and phosphatases indicates enhanced signaling regulation during maturation, whereas the marked downregulation of methyltransferases suggests attenuation of selected chromatin-modifying programs. In parallel, differential regulation of ubiquitin ligases, deubiquitinases, and other protein turnover-associated enzymes highlights proteostasis as an additional regulatory layer during neuronal differentiation. These findings provide experimentally testable entry points for defining stage-specific regulatory processes that contribute to neuronal maturation.

### Functional analysis of six transcription factors during early LUHMES neuronal differentiation

We selected six transcription factors for functional validation in cells, based on their distinctive temporal expression patterns and proteomic/phosphoproteomic profiles (Figs. 2 and 5A). MYT1 and LCOR were chosen from the continuous increase cluster due to strong upregulation at both proteome and phosphoproteome levels, with MYT1 showing the highest number of phosphosites. ISL2 (late increase) and NHLH2 (early increase) were selected as strongly upregulated factors yet without significant phosphorylation changes. DACH1, from the stable cluster, was included based on its reported role in cilia biology(Umair *et al*, 2021). Cilia have been shown to impact neuronal differentiation(Coschiera *et al*, 2023). ONECUT2, from the Day 3 valley cluster, was selected due to its highly dynamic expression pattern and known involvement in early neuronal differentiation(Sagner *et al*, 2021).

**Figure 5.**
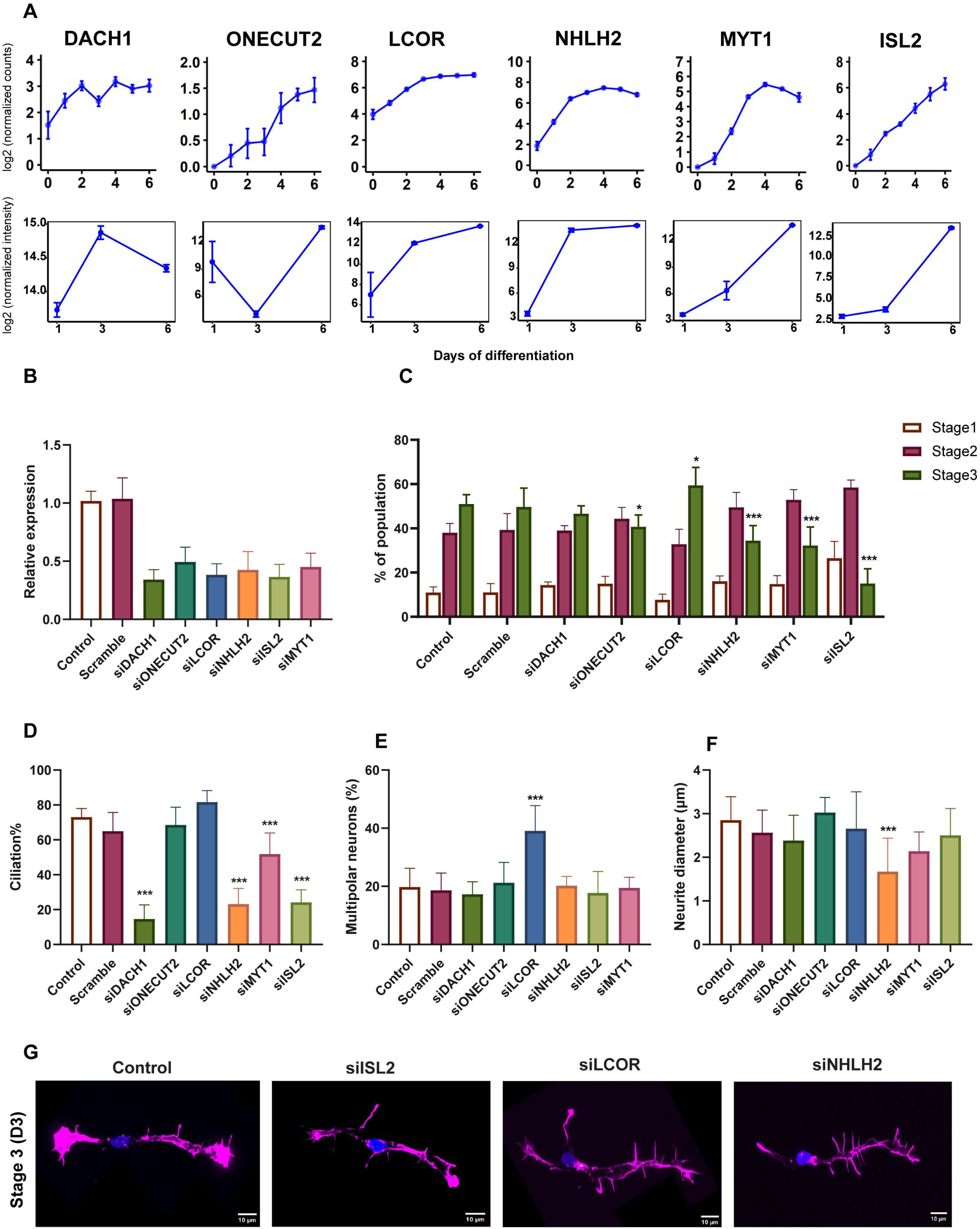
Functional validation of transcription factors during LUHMES neuronal differentiation. **(A)** Temporal expression profiles of DACH1, ONECUT2, LCOR, NHLH2, MYT1, and ISL2 during LUHMES differentiation. Upper panels show STRT-mRNA-seq expression dynamics across differentiation days (D0–D6), and lower panels show the corresponding protein abundance changes from total proteome analysis across Day 1, Day 3, and Day 6. Error bars indicate mean ± SD. **(B)** Relative mRNA expression levels following siRNA-mediated gene knockdown of the indicated transcription factors measured by qRT–PCR at Day 3. Expression values were normalized to GAPDH and are shown relative to control cells. **(C)** Quantification of neuronal differentiation stages, measured on Day 3, following transcription factor gene knockdown. Differentiating neurons were classified into Stage 1, Stage 2, or Stage 3 based on apparent neurite morphology. **(D)** Quantification of ciliated neurons on Day 3, following transcription factor gene knockdown. **(E)** Quantification of multipolar neurons on Day 3, following transcription factor gene knockdown. **(F)** Quantification of neurite diameter on Day 3, following transcription factor gene knockdown. **(G)** Representative fluorescence images of differentiating LUHMES neurons (Day 3) following siRNA-mediated gene knockdown of ISL2, LCOR, and NHLH2 compared with control cells. Scale bars, 10 μm. D Data in B–F are presented as mean ± SD from n = 6 independent biological replicates. For imaging-based quantifications in C–F, >300 cells per condition were quantified where applicable. Data in B, D, E, and F were analyzed by one-way ANOVA followed by Tukey’s multiple-comparisons test. Data in C were analyzed by two-way ANOVA followed by Tukey’s multiple-comparisons test within each differentiation stage. Statistical significance is shown for the indicated comparisons. *P < 0.05, ***P < 0.001.

Analysis of Single-cell Tagged Reverse Transcription (STRT)-mRNA-seq data across LUHMES differentiation (D0–D6) showed that all six genes were progressively upregulated(Coschiera *et al*, 2024) (Fig. 5A). To assess their functional roles, siRNA-mediated gene knockdown was performed in proliferating LUHMES progenitor cells at D−1/D0, just prior to induction of differentiation, and developmental phenotypes were analyzed on differentiation D3. Efficient knockdown was confirmed for all genes by qRT–PCR (Fig. 5B). Neuronal differentiation was evaluated at D3 using phalloidin staining to assess cell anatomies and progressive differentiation stages(Coschiera *et al*, 2024). Under control conditions, most cells reached stage 3, fewer remained at stage 2, and only a small fraction were at stage 1 (Fig. 5C). Stage 2 neurons display neurites of similar length without clear polarity, whereas the transition to stage 3 involves symmetry breaking, with one neurite elongating to form the emerging axon and establishing clear neuronal polarity(Coschiera *et al*, 2024). A shift toward detecting on D3 earlier stages was interpreted as a delay in differentiation. Consistent with delayed differentiation, knockdown of NHLH2, MYT1, and ISL2 resulted in clear differentiation delays, whereas DACH1 and ONECUT2 showed no detectable effect. In contrast, LCOR knockdown increased the proportion of cells at stage 3, indicating enhanced differentiation (Fig. 5C).

Ciliogenesis was assessed on D3 by quantifying the proportion of ciliated cells (Fig. 5D) and cilia length (data not shown). NHLH2, MYT1, and ISL2 knockdown resulted in clear ciliary phenotypes that co-occurred with differentiation delay, consistent with a link between ciliary signaling and neuronal differentiation(Coschiera *et al*, 2024). DACH1 knockdown also produced ciliary phenotypes despite the absence of a detectable differentiation delay, in line with its previously reported role in ciliogenesis and cilia function(Umair *et al*, 2021), while also indicating that ciliogenesis defects alone are not necessarily sufficient to delay differentiation. Detailed morphological analysis further revealed gene-specific effects on neurite architecture. NHLH2 knockdown resulted in reduced neurite diameter, consistent with impaired outgrowth (Fig. 5F,G). In contrast, LCOR knockdown promoted increased neurite extension and altered polarity, with a higher proportion of multipolar cells and branching structures (Fig. 5E,G), indicating that LCOR functions as a transcriptional repressor.

Together, these functional analyses in cells identify distinct roles for MYT1, ISL2, NHLH2, and LCOR during early neuronal differentiation, linking transcription factor regulation to neuronal maturation, ciliogenesis, and neurite morphogenesis in differentiating LUHMES neurons.

### Dynamics of the enhancer-promoter interactome

To investigate enhancer-mediated regulatory hierarchies during LUHMES differentiation, we integrated enhancer-promoter interaction maps with transcriptomic, proteomic and phosphoproteomic profiles across the full neuronal differentiation trajectory (D0-D6). Across all time points, 5,990 promoters interacted with 49,282 putative enhancers(Yoshihara *et al*, 2025) (Supplementary Table 9A). To identify dynamically regulated regions, we compared interaction partners of each promoter and enhancer across time points and classified each element according to its temporal interaction pattern (Supplementary Table 9B,C). Enhancers displayed substantially greater interaction dynamics than promoters, indicating extensive enhancer rewiring during neuronal differentiation (Supplementary Fig. 7A). To infer upstream regulators, we performed transcription factor (TF) motif enrichment analysis on enhancers. Motifs corresponding to 105 TFs were significantly enriched, of which 75 (71%) were detected as either differentially expressed, differentially abundant, or differentially phosphorylated across differentiation (Supplementary Table 9D). In contrast, TF motif enrichment at promoters of differentially expressed and abundant genes was comparatively limited, indicating that enhancer-mediated regulation represents a major regulatory layer during neuronal differentiation (Supplementary Table 9E, Fig. 6A).

**Figure 6.**
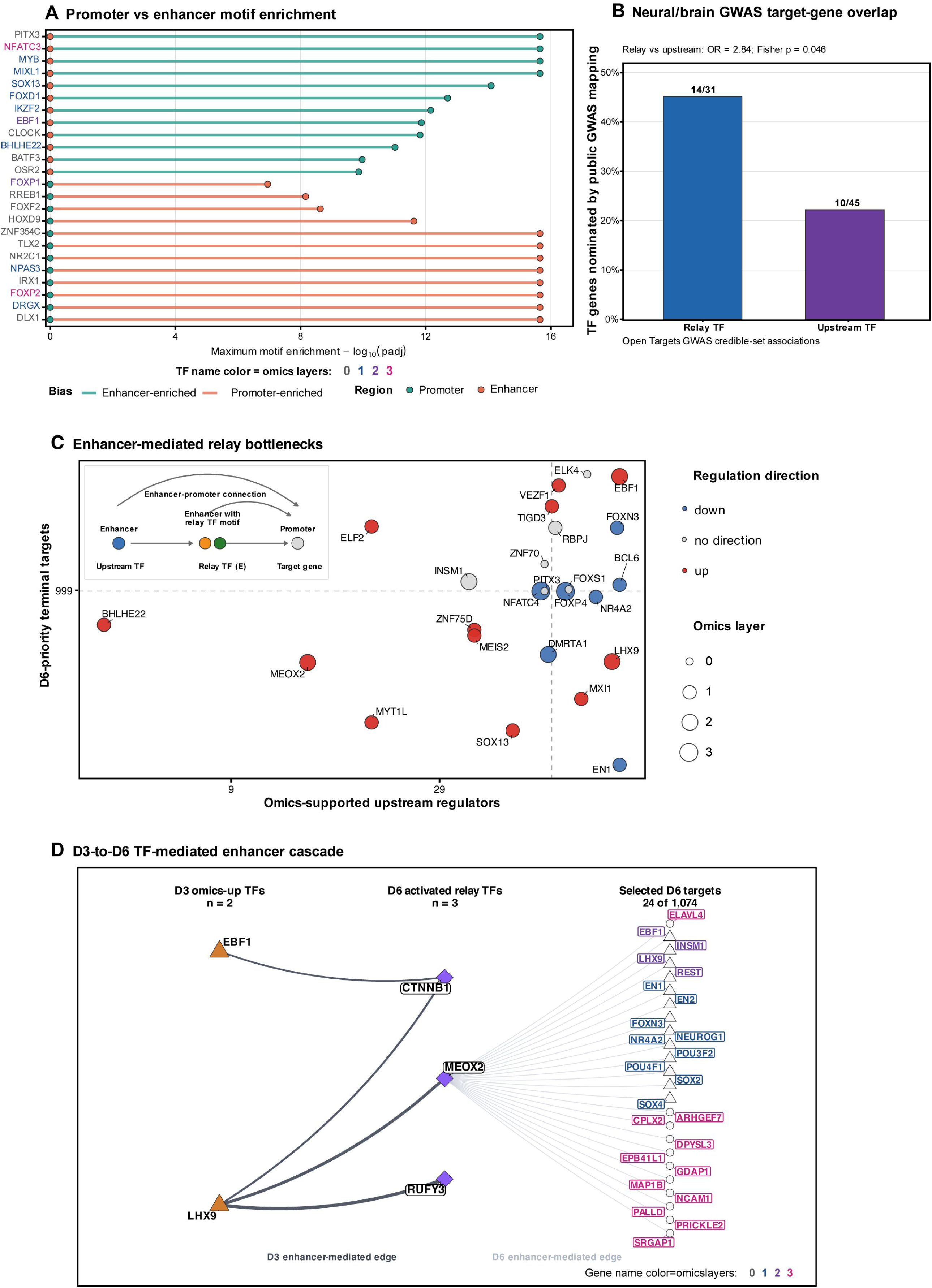
Enhancer-mediated regulatory hierarchies during LUHMES neuronal differentiation. **(A)** Comparison of TF motif enrichment in enhancer- and promoter-associated regulatory regions. TFs are ranked by maximum motif enrichment significance, and TF name color indicates the number of supporting omics layers. Lollipop stem color denotes the enrichment bias (enhancer-enriched, green; promoter-enriched, red), and dot color denotes the regulatory region in which each motif was assessed (promoter, teal; enhancer, orange). **(B)** Overlap of relay and upstream TFs with neural/brain GWAS-prioritized target genes from Open Targets credible-set associations. Relay TFs showed higher GWAS target-gene overlap than upstream TFs. **(C)** Enhancer-mediated regulatory hierarchy identifying relay TF bottlenecks. Upstream TFs were defined as TFs predicted to bind enhancers, relay TFs as enhancer-regulated TFs that also regulate downstream enhancers, and terminal targets as genes regulated through these enhancer-mediated connections. Node color indicates inferred regulation direction, and node size indicates the number of supporting omics layers. The x-axis indicates the number of omics-supported upstream regulators feeding into each TF, and the y-axis the number of D6-priority terminal target genes it regulates, so that relay TFs positioned toward the upper right act as regulatory bottlenecks; the inset schematic depicts the underlying enhancer→relay-TF→target-gene relationship. **(D)** D3-to-D6 TF-mediated enhancer cascade linking D3 omics-up TFs to D6 activated relay TFs and selected D6 target genes. Edges represent enhancer-mediated regulatory connections active at D3 or D6, highlighting late-stage regulatory programs associated with neuronal maturation. Node shapes distinguish D3 omics-up TFs, D6 activated relay TFs, and selected D6 target genes. Orange triangles indicate D3 omics-up TFs, purple diamonds indicate D6 activated relay TFs, and circles indicate selected D6 target genes. Dark and light grey edges denote D3- and D6-active enhancer-mediated connections, respectively. Gene name color in the selected D6 target layer indicates the number of supporting differential omics layers, ranging from 0 to 3.

We next constructed TF–target gene regulatory networks for each time point by linking TFs to genes if their interacting enhancers contained the corresponding TF binding motif (Methods). The resulting network comprised 7,506 nodes and 408,260 edges (Supplementary Table 9F,G). Target genes of differentially upregulated TFs at the protein level were more likely to be transcriptionally upregulated than expected by chance (OR = 2.07, p = 0.012; 2.72% vs 1.33%,

Fisher’s exact test). Similarly, target genes of significantly phosphorylated TFs were more likely to be upregulated (OR = 1.29, p = 0.0427; 18.9% vs 15.4%), supporting enhancer-mediated TF regulation during differentiation. Hierarchical reconstruction of TF cascades across time points revealed distinct regulatory layers. Several upstream regulators, including NFIA, FOXD1 and RFX7, were detected primarily at the transcriptomic level, whereas downstream regulators such as TCF3, LHX9 or ETV1 showed strong regulation at the proteomic level (Supplementary Fig. 7B). In addition, regulators including FOXN3 and PLAGL2 were identified primarily through differential phosphorylation, demonstrating the added value of integrating multi-omics modalities to resolve regulatory hierarchy (Supplementary Fig. 7B).

We next built a regulatory hierarchy using enhancer-mediated TF-target gene relationships based on their regulatory inputs. TFs that are solely predicted to bind to enhancers (i.e., only with outgoing edges) are classified as upstream TFs, whereas TFs that are themselves enhancer-regulated and predicted to bind to enhancers are classified as relay TFs (i.e., both outgoing and ingoing edges). Finally, genes only regulated by TFs are classified as terminal genes or terminal TFs, if they themselves were TFs (Fig. 6C). Most target genes were connected to enhancers containing both upstream and relay TF motifs. However, several relay TFs including ELK4, TIGD3 or EBF1 formed regulatory bottlenecks, receiving input from multiple upstream TFs while in turn regulating numerous downstream targets through their associated enhancers(Makadia *et al*, 2015) (Fig. 6C). As relay TFs emerged as bottleneck regulators within the enhancer-mediated hierarchy, we next asked whether this network position was also associated with greater relevance to human neural and brain-related genetic variation. We therefore compared public GWAS-prioritized target-gene overlap between relay and upstream TFs. Indeed, relay TFs were more likely than upstream TFs to be neural and brain-related GWAS targets (45% vs 22%; 2.84-fold; Fisher’s exact test p = 0.046), suggesting that this regulatory layer may be preferentially enriched for disease-relevant genes (Fig. 6B). To identify late-stage regulatory programs, we focused on TFs that were differentially abundant or phosphorylated at D3 and enriched in D3-active enhancers of TFs that became differentially abundant at D6 (Fig. 6D). This enabled reconstruction of enhancer-mediated TF cascades driving terminal neuronal maturation. Among these, MEOX2 emerged as a prominent late-stage regulatory hub. MEOX2 was significantly upregulated at the protein level and its binding motif was enriched in active enhancers connected to 1,074 target genes. These targets were significantly enriched for dopaminergic neurogenesis (FDR = 1.6 × 10^-2^), including key regulators and markers such as NR4A2, DDC, RET, EN1, EN2, SHH, SLC6A3, SLC18A2, and SOX2. These findings nominate MEOX2 as a previously underappreciated candidate enhancer-associated regulator of late-stage dopaminergic neuron differentiation.

## DISCUSSION

In this study, we present a comprehensive multi-omics characterization of human dopaminergic neuronal differentiation using the LUHMES cell model by integrating total proteome, phosphoproteome, transcriptome, and enhancer–promoter interactome datasets across three consecutive differentiation stages. This integrated approach reveals coordinated molecular remodeling across chromatin-associated, transcriptional, and post-translational regulatory layers, providing a systems-level view of the regulatory networks underlying human neuronal maturation.

Our proteomic profiling identified approximately 7,600 proteins per sample, with over 7,000 shared across all time points, establishing a comprehensive baseline for the LUHMES neuronal proteome. The extensive remodeling observed between Day 6 and Day 1, including 1,087 upregulated and 1,168 downregulated proteins, highlights the pronounced, large-scale transition from proliferative progenitor-like states toward differentiated neuronal phenotypes. These findings extend previous proteomic studies of LUHMES differentiation, which quantified approximately 4,000–5,000 proteins using binary comparisons between undifferentiated and differentiated states(Hao *et al*, 2026; Tüshaus *et al*, 2021), by providing increased proteome depth together with temporal resolution across defined intermediate differentiation stages. Consistent with previous reports, we observed strong upregulation of established neuronal markers including L1CAM, MAPT, SYN1, and α-synuclein, accompanied by broad repression of cell cycle and DNA replication machinery(Scholz *et al*, 2011; Tüshaus *et al*, 2021; Hao *et al*, 2026). In addition, deeper proteome coverage revealed coordinated upregulation of proteins involved in axon guidance, synaptic vesicle trafficking, and ion channel signaling, reflecting progressive acquisition of neuronal connectivity and functional maturation functions(Kolodkin & Tessier-Lavigne, 2011; Südhof, 2013; Alexander *et al*, 2015).

The phosphoproteomic component of this study represents, to our knowledge, one of the first comprehensive site-specific phosphorylation analyses during LUHMES neuronal differentiation. Across all comparisons, the phosphoproteome showed a progressive shift from phosphorylation of cell cycle regulators, chromatin-associated factors, and DNA replication machinery toward extensive modification of cytoskeletal scaffolds, synaptic components, and neuronal signaling proteins. In line with this transition, MAPT, MAP2, NEFM, DCX, and ANK3 exhibited robust increases in phosphorylation across multiple residues between Day 1 and Day 6, reflecting coordinated post-translational regulation of neuronal cytoskeletal architecture during maturation(Hao *et al*, 2026; Lyu *et al*, 2025; Yoon *et al*, 2022). These phosphorylation patterns are consistent with recent phosphoproteomic studies of iPSC-derived neuronal differentiation(Hao *et al*, 2026) and emphasize that phosphorylation functions as a complementary regulatory layer to protein abundance changes during neuronal differentiation. Consistent with these global phosphoproteome changes, kinase activity estimation indicated a shift from CDK- and PLK1-driven proliferative signaling toward MAPK7 (ERK5) and DYRK2-mediated neuronal pathways(Salaün *et al*, 2008; Wang *et al*, 2014; Santos-Durán & Barreiro-Iglesias, 2022). This kinase-level rewiring was accompanied by coordinated phosphorylation changes across transcription factor substrates, suggesting close coupling between signaling networks and transcriptional regulatory programs during differentiation. Together, these findings indicate that kinase network remodeling represents an additional regulatory layer coordinating phosphorylation dynamics during neuronal maturation.

In addition to structural and signaling proteins, we observed dynamic regulation of multiple regulatory protein classes across differentiation, including transcription factors, kinases, phosphatases, methyltransferases, and ubiquitin-associated enzymes. The increasing number of differentially regulated transcription factors toward Day 6 suggests gradual remodeling of the transcriptional landscape as cells progress toward neuronal maturation. Among transcription factor families, C2H2 zinc finger proteins were the most frequently represented across differentiation. Given the established roles of C2H2-type zinc finger proteins in brain development and neuronal specification(Al-Naama *et al*, 2020), this enrichment likely reflects broad participation of this TF family in neuronal regulatory programs. Neuronal lineage-associated factors such as MYT1 and LHX2 were upregulated, consistent with their reported roles in neuronal development(Chen *et al*, 2020; Bose *et al*, 2025; Vasconcelos & Castro, 2017). In parallel, regulatory enzyme classes also showed extensive remodeling, including increased abundance of signaling-associated kinases and phosphatases together with downregulation of several chromatin-associated methyltransferases. A subset of upregulated transcription factors, including MYT1, ST18, and MECP2, additionally showed increased phosphorylation, indicating that transcriptional regulators are coordinated across both protein abundance and post-translational regulatory layers during LUHMES differentiation.

Functional knockdown experiments further supported the importance of transcription factors identified through the integrated proteomic and phosphoproteomic analyses. Knockdown of MYT1, ISL2, and NHLH2 delayed neuronal differentiation and was accompanied by ciliogenesis-associated phenotypes, consistent with previous studies linking primary cilia function to neuronal differentiation and signaling(Coschiera *et al*, 2024). In contrast, DACH1 depletion altered ciliogenesis without producing a detectable differentiation delay, indicating that ciliary defects alone are not necessarily sufficient to impair neuronal maturation. Notably, LCOR depletion enhanced neurite extension and increased the proportion of morphologically mature neurons, suggesting that LCOR may function as a negative regulator of early neuronal differentiation.

Integration of enhancer–promoter interaction maps with transcriptomic, proteomic, and phosphoproteomic datasets provided additional insight into the regulatory architecture underlying LUHMES neuronal differentiation. Consistent with previous chromatin interaction studies(Yoshihara *et al*, 2025), enhancers displayed substantially greater interaction dynamics than promoters, supporting a model in which promoters function as relatively stable regulatory hubs while enhancers provide dynamic cell type–specific regulatory inputs during neuronal maturation. In agreement with this model, transcription factor motif enrichment was markedly stronger at dynamic enhancers than at promoters, further supporting enhancer-mediated regulation as a major regulatory layer during neuronal differentiation. Multi-omics integration further revealed coordinated regulation of transcription factors across chromatin, protein abundance, and phosphorylation layers. Several transcription factors identified through enhancer-associated motif enrichment were also differentially abundant or phosphorylated during differentiation, suggesting convergence of chromatin remodeling and post-translational regulatory mechanisms during neuronal maturation. Reconstruction of enhancer-mediated TF regulatory hierarchies additionally identified MEOX2 as a prominent late-stage regulatory hub associated with dopaminergic neurogenesis-related targets, supporting a potential role for enhancer-associated TF cascades in late neuronal maturation programs. This finding is of particular interest because MEOX2 has been studied mainly in mesodermal and vascular contexts, including Alzheimer’s disease-associated cerebrovascular dysfunction(Wu *et al*, 2005; Soto *et al*, 2016; Kokotović *et al*, 2022), whereas its role in CNS dopaminergic neuronal differentiation remains largely unexplored. More broadly, our enhancer-mediated hierarchy complements recent functional and multi-omics studies defining TF requirements in human neuronal differentiation, including a genome-wide CRISPR-Cas9 screen of approximately 1,900 TFs during NEUROG1/2-induced neurogenesis and time-resolved multi-omics analysis identifying LBX1, NHLH1, and NR2F1/2 as facilitators of human midbrain dopaminergic differentiation(Lu *et al*, 2023; Gomez Ramos *et al*, 2023). By integrating enhancer–promoter interactions with transcriptomic, proteomic, and phosphoproteomic regulation in LUHMES cells, our framework extends these studies by linking TF hierarchy to dynamic enhancer rewiring, protein abundance, and phosphorylation changes during dopaminergic neuronal maturation.

Beyond providing mechanistic insights into neuronal differentiation, our dataset has broader implications for understanding neurodegenerative diseases. Several proteins upregulated during LUHMES differentiation, including α-synuclein (SNCA), MAPT, and UCHL1, are directly implicated in Parkinson’s disease, Alzheimer’s disease, and other neurodegenerative conditions(Padilla-Godínez *et al*, 2025; Porchietto *et al*, 2025). The presence of these disease-associated proteins among differentiation-regulated targets suggests that the molecular programs driving neuronal maturation partially overlap with those disrupted in disease states. The phosphoproteomic data further complement this perspective by identifying phosphorylation events on disease-relevant proteins such as MAPT (tau), whose hyperphosphorylation is a hallmark of tauopathies(Wegmann *et al*, 2021; Watamura *et al*, 2025). LUHMES cells, with their dopaminergic identity and sensitivity to neurotoxic insults^16,17^, are particularly well suited for exploring the intersection of developmental programming and disease vulnerability. Our multi-omics resource provides a molecular framework for investigating how perturbations in the regulatory networks that drive neuronal maturation may contribute to neurodegenerative pathology.

This study has certain limitations that should be considered when interpreting the results. The LUHMES cell model, while providing excellent reproducibility and near-synchronous differentiation, represents a single neuronal subtype (dopaminergic) and may not fully capture the diversity of regulatory mechanisms operating across different neuronal lineages. The three time points sampled (Day 1, Day 3, and Day 6) provide highly informative snapshots of differentiation but may miss transient regulatory events occurring between these stages. Additionally, the phosphoproteome analysis, while comprehensive, captures predominantly serine and threonine phosphorylation due to the enrichment strategy employed; other post-translational modifications such as ubiquitination, acetylation, and methylation likely contribute additional layers of regulation during neuronal differentiation. Future studies incorporating additional time points, single-cell resolution, and broader post-translational modification coverage could further refine our understanding of the temporal dynamics underlying neuronal maturation.

In conclusion, this study provides a multi-layered resource for investigating the regulatory networks that govern human neuronal differentiation. By integrating chromatin interaction dynamics with proteomic, phosphoproteomic, and transcriptomic data, we reveal how coordinated changes across molecular layers drive the transition from proliferative progenitors to functional neurons. The identification of transcription factors regulated at multiple molecular levels, combined with dynamic enhancer–promoter rewiring, highlights the multi-layered nature of differentiation control. This dataset serves as a comprehensive reference for the LUHMES (and other) human neuronal cell models and provides a rich source of candidate regulatory factors and pathways that may be relevant to neurodevelopmental and neurodegenerative diseases.

## METHODS

### LUHMES Proliferation and Differentiation in Cell Culture

The LUHMES neuronal cell line was obtained from the American Type Culture Collection (ATCC; CRL-2927) and cultured under standard conditions (37 °C, 5% CO₂) as previously described(Lauter *et al*, 2020; Coschiera *et al*, 2023). Culture flasks were pre-coated with 50 μg/mL poly-L-ornithine hydrobromide (Sigma-Aldrich, P3655) for 1 h, followed by 1 μg/mL human fibronectin (Sigma-Aldrich, F1056) for an additional hour. After a single wash with sterile distilled water, LUHMES cells were seeded at a density of 2.5–3.5 × 10⁴ cells/cm² and maintained in DMEM/F-12 Ham growth medium (Sigma-Aldrich, D6421) supplemented with 2.5 mM L-glutamine (Sigma-Aldrich, G7513), 1× N-2 supplement (Gibco, 17502-048), and 40 ng/mL human basic fibroblast growth factor (bFGF; Thermo Fisher Scientific, PHG0369). Cells were passaged upon reaching approximately 80% confluency using either TrypLE Express (Thermo Fisher Scientific, 12605010) or 0.025% trypsin/EDTA. For neuronal differentiation, bFGF was replaced with 1 μg/mL tetracycline hydrochloride (Sigma-Aldrich, T7660) to suppress v-myc transgene expression and induce cell cycle exit. Following the addition of differentiation medium, cells were maintained in culture and harvested at Day 1, Day 3, and Day 6 to monitor neuronal differentiation progression. Prior to collection, cells were washed twice with phosphate-buffered saline (PBS), detached using TrypLE Express enzyme (Thermo Fisher Scientific, 12605010), and centrifuged at 200 × g for 5 min to remove the supernatant. Cell pellets were then snap frozen and stored at −80 °C for subsequent analyses.

### Staining and Microscopy

Staining and microscopy analyses were performed as previously described(Coschiera *et al*, 2024) with some modifications. Briefly, 2.5×10^4^ LUHMES cells were seeded onto pre-coated cover glasses in 12-well plates 24 hours prior to differentiation. Cells were fixed with 2% PFA for 5 minutes and permeabilized with 0.2% Triton X-100 in PBS for 12 minutes. After blocking with 2% BSA and 0.1% Tween-20 in PBS for 30 minutes, coverslips were incubated with Alexa Fluor 488 Phalloidin (Thermo Fisher Scientific, A12379) overnight at 4℃ and nuclei were counterstained with Hoechst 33342. Coverslips were mounted with ProLong Glass antifade mountant and cured in the dark at room temperature for 24 hours to achieve optimal refractive index. Confocal images were acquired using a Nikon Ti2 inverted microscope equipped with a Crest Optics X-Light V3 spinning disk unit and a Kinetix sCMOS camera.

### Total Proteome and Phosphoproteome Sample Preparation

For total proteome and phosphoproteome analyses, three independent biological replicates were analyzed for each differentiation stage. LUHMES cell pellets (2 x 10^6^ cells/replicate) were lysed in 8 M urea prepared in 50 mM ammonium bicarbonate (AMBIC; Sigma-Aldrich), supplemented with phosphatase and protease inhibitor cocktails (Sigma-Aldrich, P2745 and P8340). Lysis was performed on ice, followed by two rounds of sonication in a water bath for 15 min each, with a 10 min incubation on ice in between. Lysates were centrifuged at 16,000 × g for 20 min at 4 °C, and the supernatants were collected for downstream processing. Total protein concentration was determined using a BCA protein assay (Thermo Fisher Scientific), and aliquots were adjusted to reduce urea concentration for accurate measurement. Equal protein amounts (250 µg) were reduced with tris(2-carboxyethyl) phosphine (TCEP; Sigma-Aldrich) and alkylated with iodoacetamide. Samples were diluted 4-fold with 50 mM AMBIC to reduce urea concentration below 2 M and digested overnight at 37 °C with sequencing-grade modified trypsin (Promega) at a 1:50 to 1:20 enzyme-to-protein ratio. The resulting peptides were desalted using C18 Macrospin columns (Nest Group) and split into two portions: one for total proteome analysis and one for phosphopeptide enrichment. For phosphoproteome preparation, peptides were loaded onto stage tips packed with Ti⁴⁺-IMAC microspheres pre-equilibrated with loading buffer (80% acetonitrile (ACN), 6% trifluoroacetic acid (TFA)). After binding, nonspecifically bound peptides were removed by sequential washes with wash buffer I (50% ACN, 6% TFA, 200 mM NaCl) and wash buffer II (50% ACN, 0.1% TFA). Phosphopeptides were eluted using 10% ammonium hydroxide, and eluates were lyophilized prior to LC-MS/MS analysis.

### LC-MS/MS analysis

Desalted peptide samples were analyzed using the Evosep One liquid chromatography system coupled to a timsTOF Pro 2 hybrid trapped ion mobility quadrupole time-of-flight mass spectrometer (Bruker Daltonics) via a CaptiveSpray nano-electrospray ion source. Peptide separation was performed on an 8 cm × 150 µm column packed with 1.5 µm C18 beads (EV1109, Evosep) using the 60 samples-per-day workflow, which employed a 21-minute gradient. Mobile phases consisted of 0.1% formic acid in water (solvent A) and 0.1% formic acid in ACN (solvent B). Data-independent acquisition (DIA) was conducted in positive-ion mode using the DIA-PASEF acquisition strategy with sample-specific optimized parameters. To guide DIA-PASEF optimization, data-dependent acquisition (DDA) runs in PASEF mode were first performed on pooled samples. DIA-PASEF acquisition settings were then refined for each sample type using timsControl software (Bruker Daltonics). For phosphoproteome samples, the DIA-PASEF parameters included a mass range of 433–1684 m/z, mobility range of 0.85–1.30 1/K₀, and an estimated cycle time of 1.80 s. For total proteome analysis, the mass range was set to 370–1370 m/z, with the same mobility range and an estimated cycle time of 1.27 s. Ion mobility windows were defined based on the ion cloud distribution obtained from the corresponding DDA runs. Following acquisition, raw data from both total proteome and phosphoproteome experiments were processed using DIA-NN software (version 1.8.1). For total proteome analysis, search parameters included a maximum of one missed cleavage, up to three variable modifications, N-terminal methionine excision, fixed carbamidomethylation of cysteine, variable oxidation of methionine, and acetylation at the protein N-terminus. Peptide lengths were restricted to 7–30 amino acids, with precursor charges of 2–4, a precursor m/z range of 300–1600, and fragment ion m/z range of 100–1700. Both precursor and MS1 mass accuracies were set to 15 ppm. For phosphoproteomics, DIA-NN settings were similar, except methionine oxidation was excluded to avoid confounding modification overlap with phosphorylation.

### Total and Phosphoproteome Data Analysis

Quantitative analysis of total proteome and phosphoproteome data was performed using the MSstats and MSstatsPTM R packages, respectively(Kohler *et al*, 2023b, 2023a). For total proteome analysis, precursor-level intensities were log-transformed, normalized, and summarized to protein-level abundances, including Tukey’s median polish summarization. Pairwise differential abundance testing between Day 3 vs Day 1, Day 6 vs Day 3, and Day 6 vs Day 1 was performed using linear mixed-effects model per protein. Proteins with adjusted p-value < 0.05 and |log₂(fold change)| > 1 were considered significantly differentially abundant. For phosphosite analysis, phosphopeptide-level intensities from DIA-NN were processed using MSstatsPTM and differential phosphorylation between conditions was assessed. Phosphosites with an adjusted p-value < 0.05 and |log₂(fold change)| > 1 were considered significantly regulated.

### Gene Ontology and Pathway Enrichment Analysis

To investigate the functional changes associated with neuronal differentiation, we performed Gene Ontology (GO) enrichment analysis on differentially abundant proteins using the DAVID Knowledgebase v2025(Huang *et al*, 2009; Sherman *et al*, 2022). Proteins showing significant changes in abundance were identified from pairwise comparisons between Day 1 vs Day 3, Day 3 vs Day 6, and Day 1 vs Day 6 of differentiation. Enrichment analysis was conducted across the three GO categories: Biological Process (BP), Cellular Component (CC), and Molecular Function (MF). Additionally, Reactome pathway analysis(Ragueneau *et al*, 2026) was applied to identify signaling and metabolic pathways associated with differentiation-related protein dynamics. These analyses provided insights into the molecular processes and pathways that underlie the progression of neuronal differentiation in LUHMES cells.

### Enhancer-Promoter Interactome Analysis

Chromatin interaction data for LUHMES cells were obtained from previous publication(Yoshihara *et al*, 2025) and converted to hg38 genome coordinates using liftOver tool available via Bioconductor package in R. All converted interaction coordinates, including promoter–enhancer pairings, are available in Supplementary Table 9A.

### Rewiring index

Dynamic classification of promoter–enhancer interactions across differentiation was performed based on the presence or absence of each interaction at Day 1 (D1), Day 3 (D3), and Day 6 (D6). Interacting elements were assigned to the following categories: **Gained_early:** absent at D1, present at D3 and D6, **Lost_early:** present at D1, absent at D3 and D6, **Lost_late:** present at D1 and D3, absent at D6, **Gained_late:** absent at D1 and D3, present at D6, **Stable:** present at D1, D3, and D6, **Other_pattern:** any other combination.

For each promoter or enhancer element, their interacting partners were listed at each time point. For each promoter, partner enhancer sets were enumerated at each timepoint and Jaccard similarity was computed for adjacent transitions (D1–D3, D3–D6). The rewiring index was defined as:

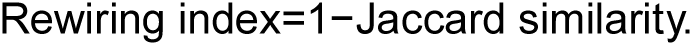

Dynamic class-wise summaries used the biologically relevant transition for each category (early classes: D1–D3; late classes: D3–D6; Stable: mean of both). The promoter and enhancer classifications are given in Supplementary Table 9B,C.

### Motif Scanning and enrichment analysis

Transcription factor motifs are retrieved from JASPAR 2024 (using CORE Homo sapiens subset) via TFBSTools R package. Position weight matrices (PWMs) were converted using toPWM(type = “log2probratio”, pseudocounts = 0.8). Motif names and family annotations were extracted from TFBSTools metadata. For timepoint-specific analyses, foreground consisted of dynamically changing promoters or enhancers for a given comparison and background consisted of stable promoters or enhancers. We computed GC content and width per element and used them as covariates in enrichment modeling to control sequence composition and length. Motif enrichment was estimated by fitting, per motif, a binomial GLM contrasting foreground (y=1) vs background (y=0):

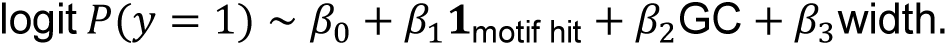

The coefficient *β*_1_ and its standard error were extracted (fit_one), yielding odds ratios (OR), Wald p-values, and Benjamini–Hochberg FDR (padj) per motif. Motifs were considered enriched if coef > 0 and padj < 1×10⁻³. Supplementary Table 9D,E lists all the enriched TFs in dynamic enhancers and promoters, respectively.

### Public GWAS target-gene overlap of upstream and relay TFs

To assess whether transcription factors occupying different positions in the enhancer-mediated regulatory hierarchy differed in their connection to human genetic associations, we compared public GWAS-prioritized target-gene overlap between upstream and relay TFs. TF hierarchy roles were taken from the annotated TF hierarchy network, in which TF genes were classified as upstream TFs or relay TFs based on their regulatory position in the enhancer-mediated TF-target network (Fig. 6C). The analysis was performed at the level of the TF genes themselves, rather than their downstream targets, to avoid confounding from the substantial overlap between genes regulated by upstream and relay TFs. Public GWAS gene associations were obtained from Open Targets Platform target-disease associations restricted to the gwas_credible_sets data source. For each TF gene in the hierarchy, Open Targets associations were retrieved and filtered to retain a broad set of neural and brain-related traits. Trait filtering was based on disease/trait labels containing terms related to brain biology, cognition, neurodevelopmental, neurological, neurodegenerative, psychiatric, and selected brain-relevant disorders, including schizophrenia, autism spectrum disorder, attention deficit hyperactivity disorder, bipolar disorder, depressive disorders, epilepsy/seizure phenotypes, Parkinson disease, Alzheimer disease, and dementia. Broad labels likely to capture non-neural brain pathologies were excluded, including brain cancer/neoplasm, brain aneurysm, brain compression, and intracranial hemorrhage. A TF gene was considered a public neural/brain-related GWAS target if it had at least one retained Open Targets GWAS credible-set association after trait filtering. The proportion of GWAS-associated TF genes was then compared between relay TFs and upstream TFs. Enrichment was tested using a two-sided Fisher’s exact test on a 2 x 2 contingency table of TF hierarchy role and presence or absence of a retained GWAS target-gene association. The relay-versus-upstream odds ratio and Fisher’s exact test p-value were reported. As a background comparison, the same analysis was also run using all Open Targets GWAS credible-set associations without neural/brain-related trait filtering.

### siRNA Transfections

Transfections were performed by using Lipofectamine RNAiMAX (Thermo Fisher Scientific, 13778075) according to the manufacturer’s instructions(Coschiera *et al*, 2024). Briefly, LUHMES cells were seeded into 12-well plates at a density of 5×104 cells/well. On the next day, cells were transfected with negative control siRNA or target gene-specific siRNAs (Supplementary Table 8). All siRNAs were used at a final concentration of 50 nM, and cells were incubated with the siRNA–lipid complexes for 24 hours before the culture medium was replaced with differentiation medium. Following culturing for 72 hours at 37°C, transfection efficiency was assessed by qRT-PCR.

### RNA Isolation and qRT-PCR

Total RNA was extracted by using RNA Extraction Kits (Qiagen 74104) following the manufacturer’s protocol. RNA purity was assessed with a Nanodrop ND-100 (Thermo Fisher Scientific). qRT-PCR was performed with the 7500 Fast Real-Time PCR platform and the FastStart Universal SYBR Green Master Mix (Roche 4913850001). Relative mRNA levels were evaluated using the 2−ΔΔCt method and GAPDH was used as the housekeeping gene to normalize the values.(Coschiera *et al*, 2024)

## DATA AVAILABILITY

The raw mass spectrometry file names for all samples analyzed in this study (total proteome and phosphoproteome) are provided in Supplementary Table 10. All raw mass spectrometry data have been deposited in the MassIVE repository (https://massive.ucsd.edu) under accession number MSV000102026.

## AUTHOR CONTRIBUTIONS

D.M.P. and X.L. performed proteomics and phosphoproteomics experiments. H.L. and A.C. performed all LUHMES cell culture, immunocytochemistry, microscopy, and imaging work. H.L. conducted the siRNA gene knockdown experiments. Y.J. contributed to bioinformatic analyses and figure preparation. The enhancer data were obtained by M.Y. and J.K. P.Sa. integrated enhancer–promoter interaction data and rest of the modalities, performed analyses and interpretations. D.M.P. analyzed the data, prepared the figures, integrated the different datasets, and wrote the manuscript with input from all authors. The project concept rooted in tight collaboration between the groups of M.V., J.K., P.Sa., P.Sw. and M.Y., who also together supervised the study and secured funding for their laboratories. All authors reviewed and approved the final manuscript.

## ACKNOWLEDGEMENTS

D.M.P. was partially supported by the iCANPOD postdoctoral program, and Y.J. is an iCANDOC doctoral student, both funded by the iCANDOC doctoral education pilot in precision cancer medicine. This research was funded by Academy of Finland (nos. 288475 and 294173; M.V.), Sigrid Jusélius Foundation (M.V., J.K.), Finnish Cancer Foundation (M.V.), Biocenter Finland (M.V.), HiLIFE (M.V.), Swedish Research Council (P.Sw., J.K.,P.Sa.), Carl Tryggers Foundation (P.Sa.), Swedish Brain Foundation (P.Sw., J.K.), Olle Engkvist Foundation (P.Sw.), Åhlén Foundation (P.Sw.), Karolinska Institutet PhD student Fellowship (A.C., H.L.), and Chinese Scholarship Council PhD student Fellowship (H.L.). We also thank Akshara Urs for providing technical assistance during the early stages of the laboratory work.

## COMPETING INTERESTS

The authors declare no competing interests.

## SUPPLEMENTARY FIGURES

**Supplementary Figure 1.**
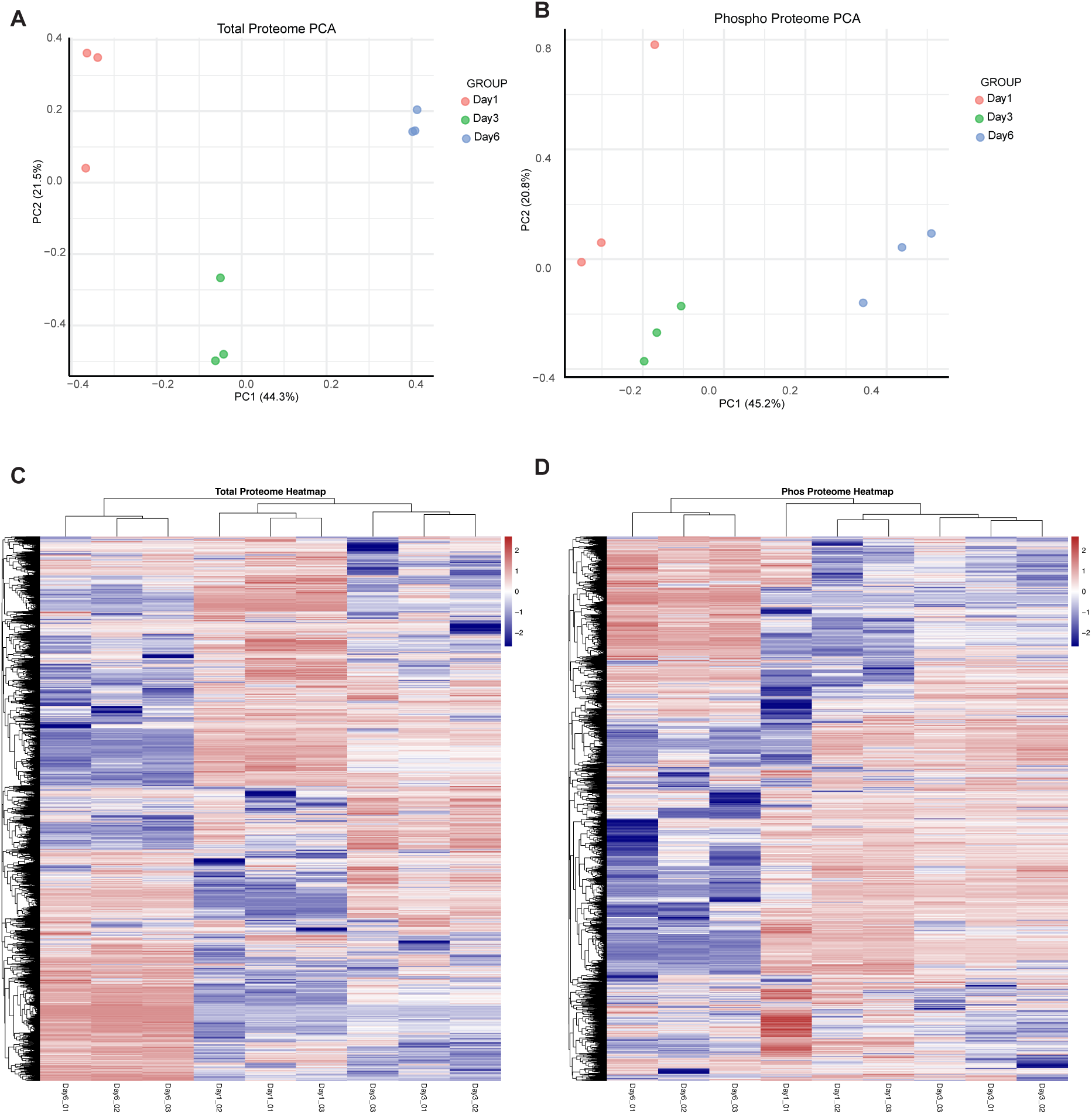
Global proteome and phosphoproteome sample clustering. **(A)** Principal component analysis (PCA) of total proteome profiles from differentiating LUHMES neurons collected at Day 1, Day 3, and Day 6. **(B)** Principal component analysis (PCA) of phosphoproteome profiles from these LUHMES neurons collected at Day 1, Day 3, and Day 6. **(C)** Heatmap showing hierarchical clustering of total proteome abundances across biological replicates from Day 1, Day 3, and Day 6. **(D)** Heatmap showing hierarchical clustering of phosphoproteome abundances across biological replicates from Day 1, Day 3, and Day 6.

**Supplementary Figure 2.**
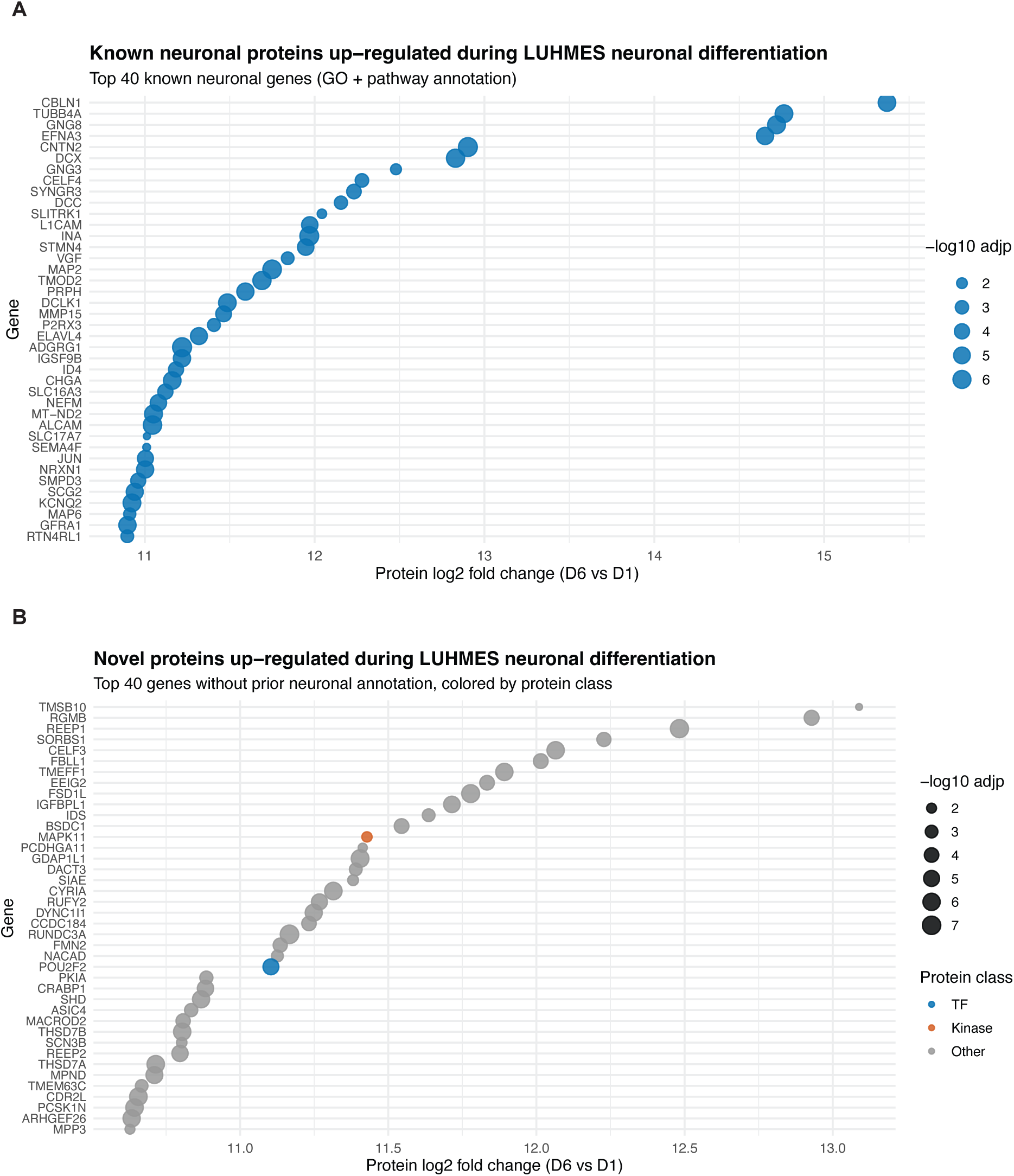
Upregulated neuronal and unannotated proteins during LUHMES differentiation. **(A)** Dot plot showing the top 40 proteins upregulated at Day 6 compared to Day 1 with known neuronal annotation based on Gene Ontology and pathway databases. Proteins are plotted by log_2_ fold change (x-axis), and dot size indicates –log_10_ (adjusted p-value). **(B)** Dot plot showing the top 40 proteins upregulated at Day 6 compared to Day 1 without prior neuronal annotation. Points are colored by protein class (transcription factors, kinases, or other), and dot size indicates –log_10_ (adjusted p-value).

**Supplementary Figure 3.**
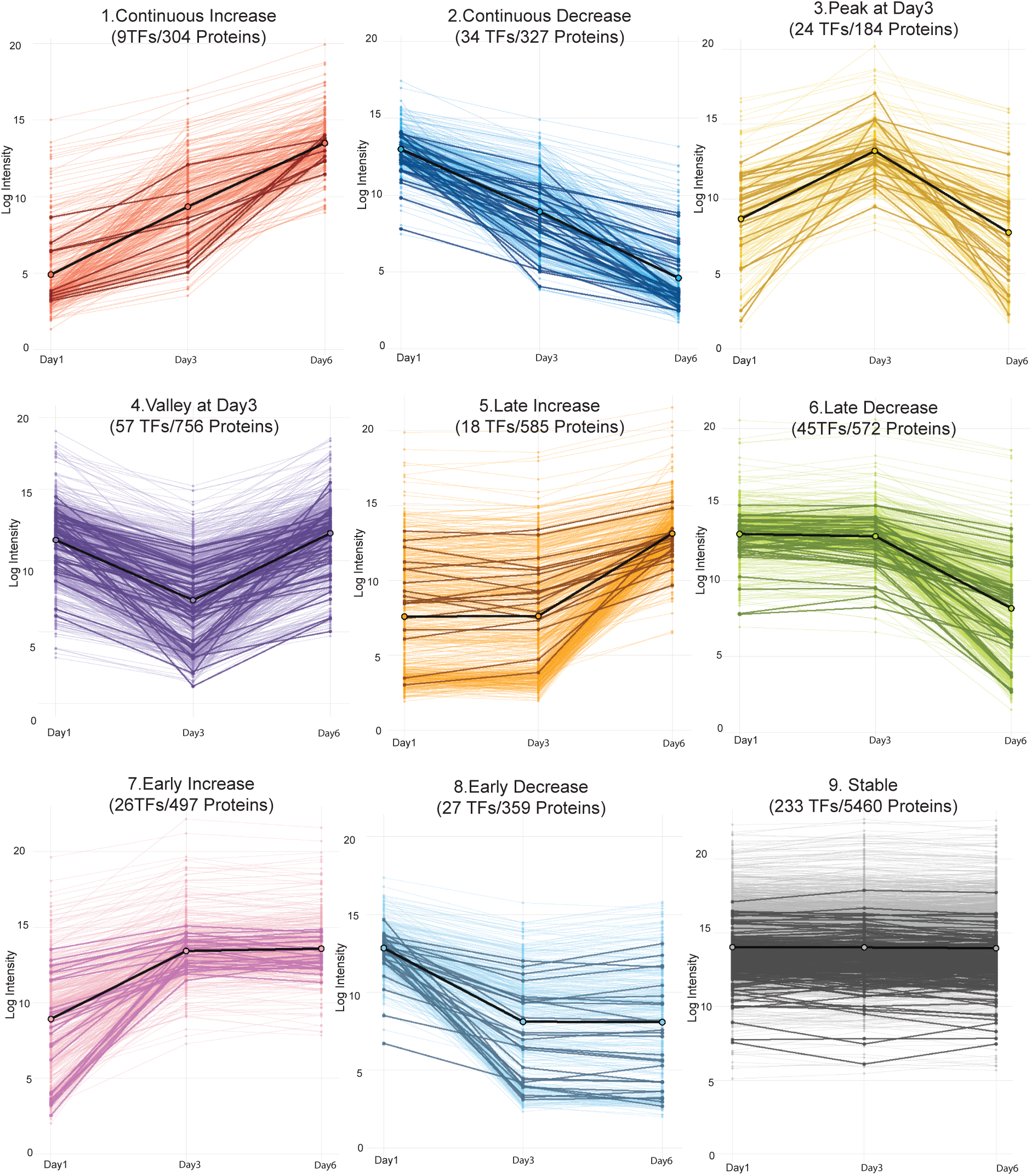
Extended clustering of protein expression trajectories during LUHMES differentiation. Line plots show protein abundance profiles across Day 1, Day 3, and Day 6, grouped into nine clusters based on their temporal expression patterns: continuous increase (1), continuous decrease (2), peak at Day 3 (3), valley at Day 3 (4), late increase (5), late decrease (6), early increase (7), early decrease (8), and stable expression (9). Each thin line represents an individual protein. Bold colored lines indicate transcription factors (TFs), and the bold black line represents the average expression profile of the cluster. The number of TFs and total proteins assigned to each cluster is indicated in each panel.

**Supplementary Figure 4.**
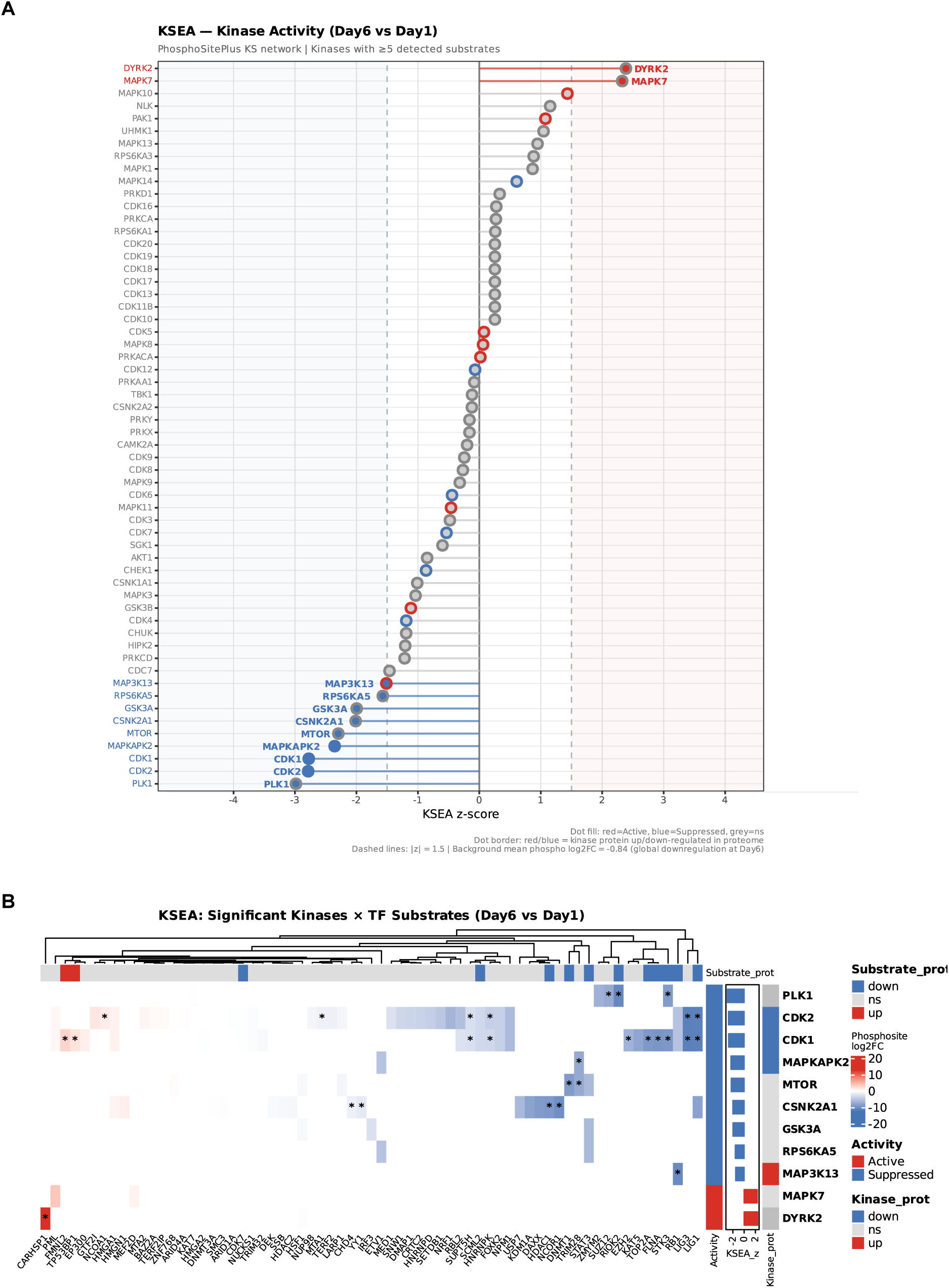
Kinase activity inference from LUHMES phosphoproteomics (Day 6 vs Day 1). **(A)** Kinase activity lollipop plot. KSEA z-scores for all 58 kinases with ≥5 detected substrates in the Day6vsDay1 phosphoproteome comparison. Each dot represents one kinase; the horizontal position indicates the z-score (positive = inferred active, negative = inferred suppressed relative to the global phosphoproteome background). Kinases meeting the significance threshold (|z| ≥ 1.5 or p < 0.05, two-tailed) are highlighted. Dot color indicates inferred activity direction (red = active, blue = suppressed). **(B)** Kinase–TF substrate heatmap. Log2 fold-change (Day6 vs Day1) of 67 transcription factor (TF) substrates phosphorylated by the 11 significant kinases. Rows represent individual TF substrates; columns represent kinases ordered by z-score (most suppressed to most active, left to right). Color scale: red = increased phosphorylation, blue = decreased phosphorylation. Asterisks (*) denote phosphosites meeting significance criteria (|log2FC| ≥ 1 and adj. p < 0.05).

**Supplementary Figure 5.**
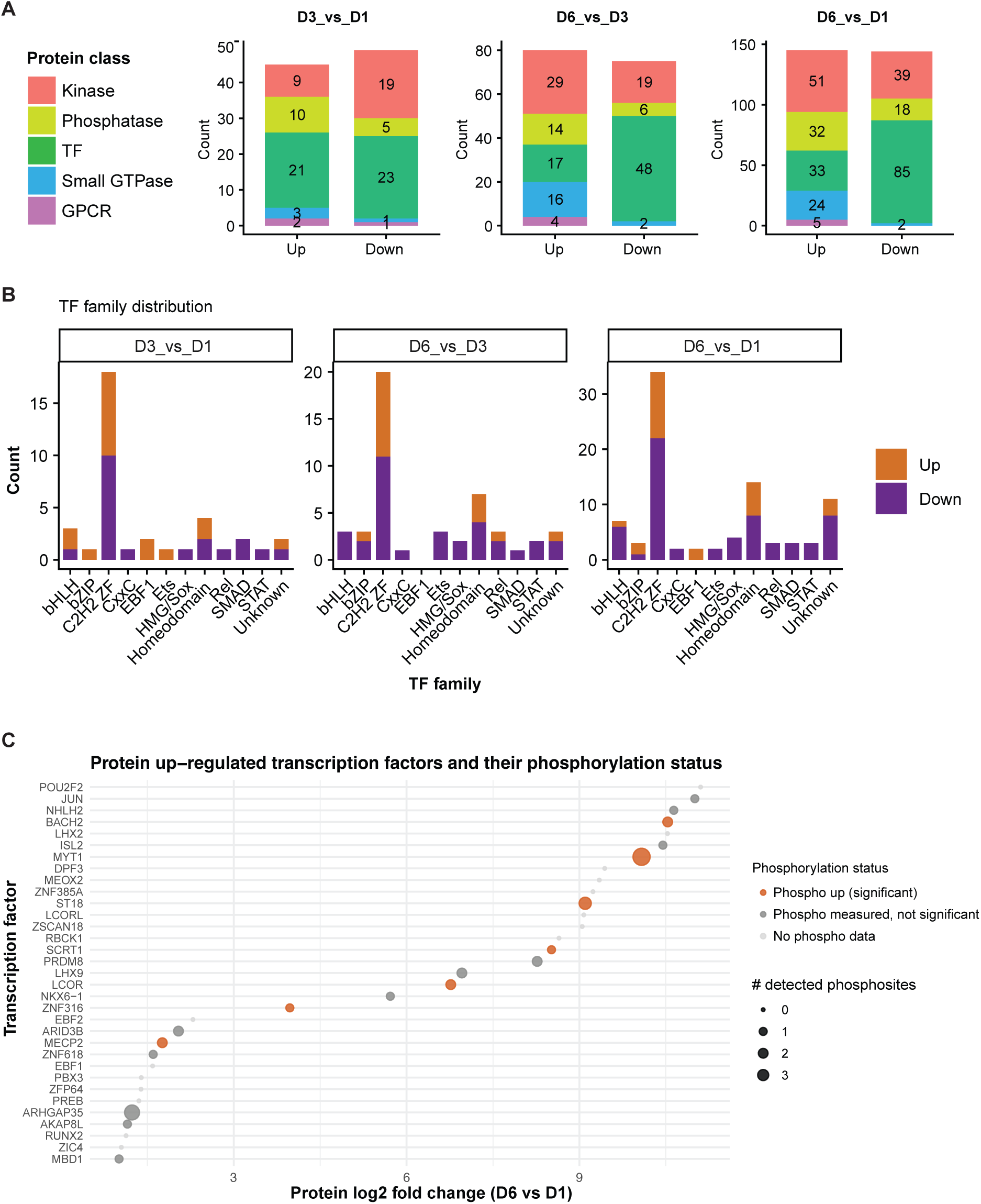
Global protein class dynamics and phosphorylation status of transcription factors during LUHMES neuronal differentiation. **(A)** Stacked bar plots showing the number of significantly upregulated and downregulated proteins across major protein classes, including kinases, phosphatases, transcription factors (TFs), small GTPases, and GPCRs, for the indicated pairwise comparisons (Day 3 vs Day 1, Day 6 vs Day 3, and Day 6 vs Day 1). **(B)** Bar plots showing the distribution of transcription factor families among significantly upregulated and downregulated TFs for each comparison (Day 3 vs Day 1, Day 6 vs Day 3, and Day 6 vs Day 1). **(C)** Dot plot showing transcription factors that are significantly upregulated at the protein level in Day 6 compared to Day 1. Each dot represents one TF plotted by its protein log_2_ fold change (x-axis). Dot size indicates the number of detected phosphosites for each TF, and dot color denotes phosphorylation status (significantly increased phosphorylation, phosphorylation measured but not significant, or no phosphoproteomic data).

**Supplementary Figure 6.**
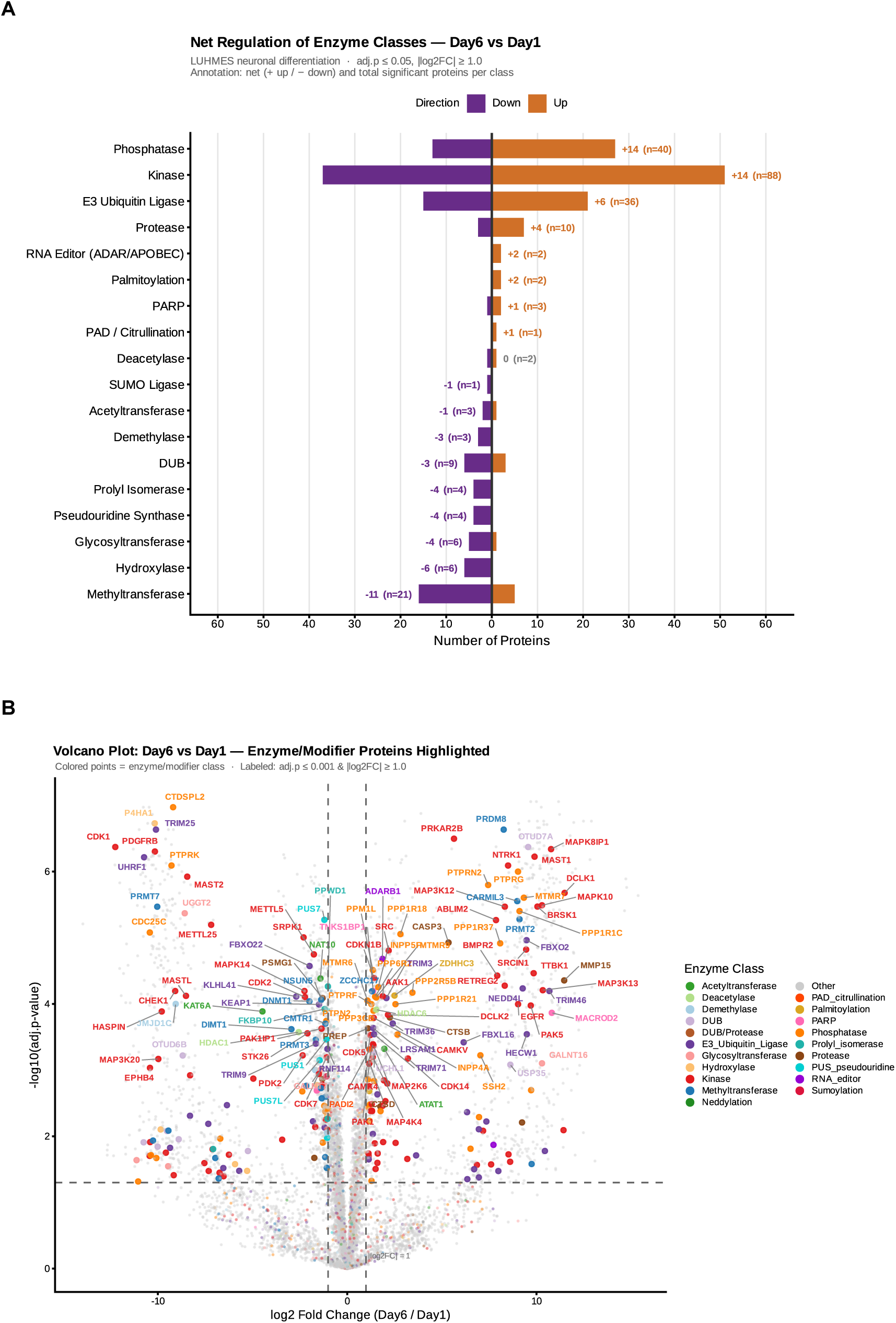
Regulation of enzyme/modifier classes during LUHMES neuronal differentiation. **(A)** Net regulation of enzyme/modifier classes at Day 6 vs. Day 1. Bars show significantly upregulated (orange) and downregulated (purple) proteins per class (adj. p ≤ 0.05, |log2FC| ≥ 1.0). Numbers indicate net direction score and total significant proteins (n). Kinases and phosphatases showed net upregulation (+14 each), whereas methyltransferases showed the strongest net downregulation (−11). **(B)** Volcano plot of enzyme/modifier proteins at Day 6 vs. Day 1. Colored points indicate classified enzyme/modifier proteins by class; grey points represent other proteins. Dashed lines indicate |log2FC| = 1.0 and adj. p = 0.05. Gene labels mark proteins with adj. p ≤ 0.001 and |log2FC| ≥ 1.0. Fold changes represent Day 6 relative to Day 1 of LUHMES dopaminergic neuronal differentiation.

**Supplementary Figure 7.**
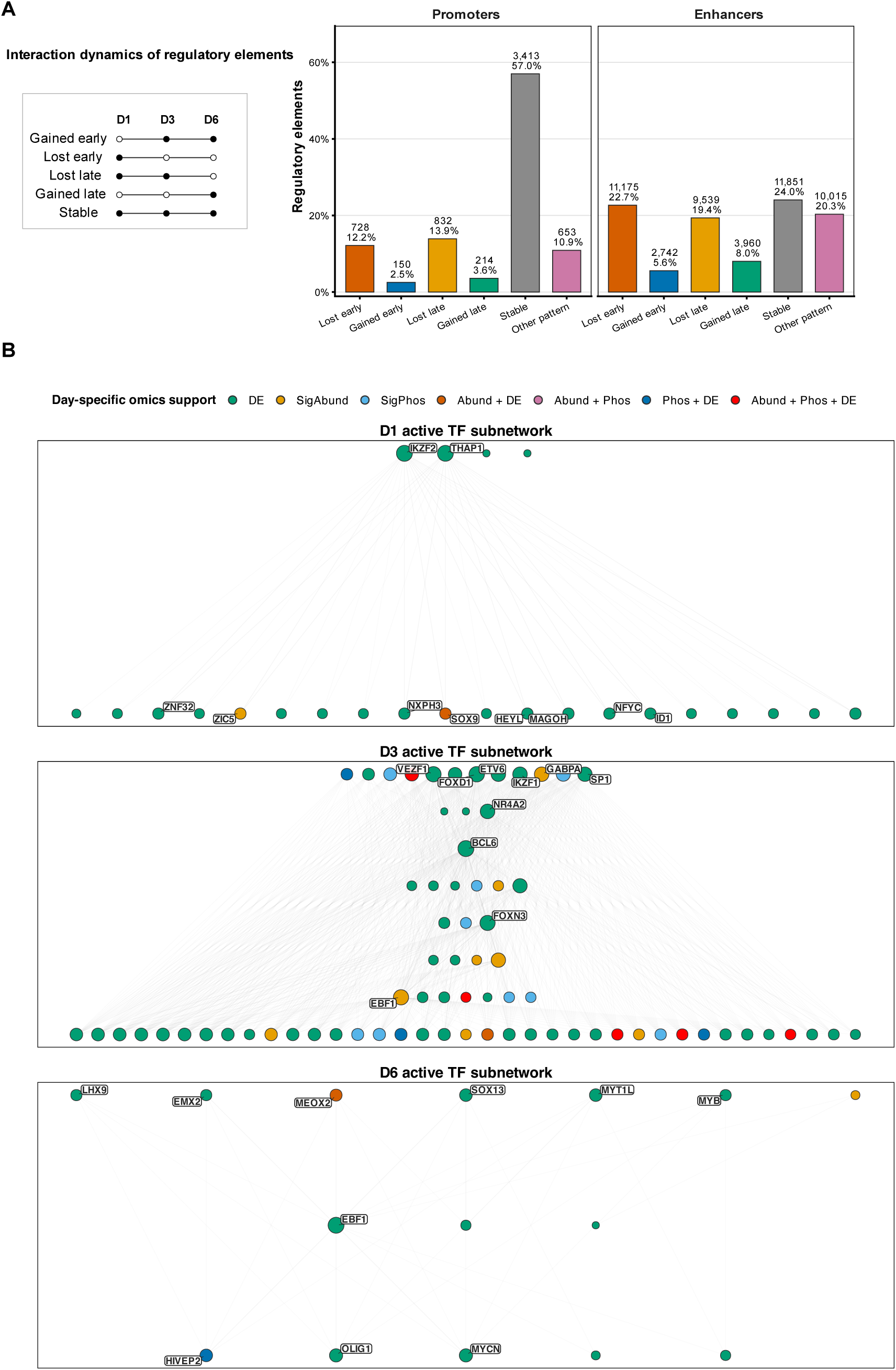
Enhancer–promoter interaction dynamics and day-specific enhancer-mediated TF subnetworks during LUHMES neuronal differentiation.(A) Promoters and enhancers were grouped according to the presence or absence of their chromatin interactions across D1, D3, and D6. Interaction patterns were classified as lost early, gained early, lost late, gained late, stable, or other pattern, as illustrated in the schematic. Bar plots show the number and percentage of promoters and enhancers in each category, highlighting that enhancers display more dynamic interaction changes, whereas promoters are more frequently stable. **(B)** Day-specific transcription factor subnetworks inferred from dynamically interacting enhancers at D1, D3, and D6. Nodes represent TFs with day-specific omics support, and edges indicate enhancer-mediated regulatory connections between TFs. Node color denotes the supporting omics modality, including differential expression, protein abundance, phosphorylation, or their combinations, while node size reflects connectivity within each subnetwork. Selected highly connected TFs are labeled.

